# Multi-species community platform for comparative neuroscience in teleost fish

**DOI:** 10.1101/2024.02.14.580400

**Authors:** Sumit Vohra, Kristian Herrera, Tinatini Tavhelidse-Suck, Joachim Wittbrodt, Simon Knoblich, Ali Seleit, Alexander Aulehla, Jonathan Boulanger-Weill, Sydney Chambule, Ariel Aspiras, Cristina Santoriello, Mark Fishman, Hans-Christian Hege, Daniel Baum, Florian Engert, Yasuko Isoe

## Abstract

Studying neural mechanisms in complementary model organisms from different ecological niches in the same animal class can leverage the comparative brain analysis at the cellular level. To advance such a direction, we developed a unified brain atlas platform and specialized tools that allowed us to quantitatively compare neural structures in two teleost larvae, medaka (*Oryzias latipes*) and zebrafish (*Danio rerio*). Leveraging this quantitative approach we found that most brain regions are similar but some subpopulations are unique in each species. Specifically, we confirmed the existence of a clear dorsal pallial region in the telencephalon in medaka lacking in zebrafish. Further, our approach allows for extraction of differentially expressed genes in both species, and for quantitative comparison of neural activity at cellular resolution. The web-based and interactive nature of this atlas platform will facilitate the teleost community’s research and its easy extensibility will encourage contributions to its continuous expansion.

## Main

Although the basic blueprint of the brain is well-conserved across vertebrates, different species exhibit diversity of cerebral subregions even in the same animal class. Using different animal models in the same class is therefore a powerful and efficient way to understand how neural circuits give rise to behavior, since it allows to uncover generally conserved, as well as individually specific mechanisms of neural function. For example, in rodent models, mountain and prairie voles are easily distinguished with respect to different mating-partnership, where oxytocinergic neurons have been identified to play a critical role in this process (Ross et al. 2009). In primate models, macaques, marmosets and, indeed, humans are used to compare the performance in recognition tasks (Miller et al. 2016; Wilson et al. 2013). However, in most of these animal models their comparatively large brains make it difficult to investigate the concerted activity brain-wide and at the single-cell level. Here we present a multi-species community platform for teleost fish that has several advantages in comparing brains across different species.

Teleost larvae are useful model organisms to elucidate behaviors and the underlying neural circuits for several reasons: (1) basic brain circuits are conserved among vertebrates (O’Connell and Hofmann 2012); (2) the small body size makes it easy to perform brain wide imaging across different sensory modalities (Herrera et al. 2021; Migault et al. 2018); and (3) the transparent body allows us to analyze the brain activity *in vivo* across the whole brain at cellular level (Ahrens et al. 2012). Zebrafish (*Danio rerio*) larvae have been therefore investigated intensely in the context of system neuroscience, but this approach has yet only sparsely been extended to different teleost species (Lloyd et al. 2022), an endeavor necessary to study behaviors and neural circuits in an explicitly comparative fashion.

Here, we selected medaka fish (*Oryzias latipes*) as another fish model species for which molecular genetic tools and genomic research are available (Kasahara et al. 2007). We chose medaka because it is phylogenetically distant from zebrafish (diverged 250 Million years ago) and inhabits a different natural environment (Furutani-Seiki and Wittbrodt 2004). Zebrafish inhabit murky freshwater, whereas medaka live in clear water and, further, have much higher salinity tolerance (Inoue and Takei 2002). This allows us to study the evolutionary diversity that emerged from different ecological niches. Although there are several brain atlases for adult (Anken and Bourrat 1998; Ishikawa, Yoshimoto, and Ito, n.d.) and larval medaka fish (Ishikawa, Yamamoto, and Hagio 2022), most previous atlases do not cover the whole brain with fine-enough spatial resolution for quantitative analysis, and region names are often not the same in the zebrafish and medaka atlases; this lack of an integrated cross-species platform makes it difficult to conduct quantitative comparative studies across teleost fish.

To address this issue, we extended the existing FishExplorer platform (https://fishexplorer.zib.de) containing the larval zebrafish brain (Z-brain) atlas (https://fishexplorer.zib.de/zebrafish/lm) that has been established and used to study neural circuits (Randlett et al. 2015; Vohra et al. 2023), to allow for the seamless incorporation of a larval medaka brain (M-brain). To compare brain anatomical regions among different fish species, we parceled each brain into the corresponding *mutually exclusive and comprehensively exhaustive* (MECE) regions, which cover the whole brain without overlaps and without gaps. This approach helps to quantitatively compare the whole brain across species in terms of volume and gene expression. In addition, neural subpopulations that express specific neurotransmitters were labeled and miscellaneous (MISC) masks were created, complementary to the MECE masks, to facilitate further comparison. Using this new and multi-species atlas-based platform, we found several different neural subclusters in the diencephalon and telencephalon that are interesting for future investigations. The interactivity and extensibility of the atlas platform makes it ideally suited as a community platform that incorporates further teleost data from different research labs.

## Results

### Establishment of multi-species atlas platform

We first needed to select zebrafish and medaka fish larvae at comparable stages of development. Although zebrafish take less time to hatch (2-3 days) than medaka (7-8 days), zebrafish do not swim regularly until 2 days after hatching, whereas medaka start swimming immediately after hatching. Also, brain and other organ development are grossly comparable at these stages (Furutani-Seiki and Wittbrodt 2004). Therefore, for all further comparison, we selected larvae 5-7 days post fertilization (dpf) of zebrafish and 7-8 dpf of medaka (Figure 1a). This is aligned with most behavioral research in zebrafish larvae, which is usually conducted at 5-7 dpf (Oteiza et al. 2017; X. Chen and Engert 2014; Kubo et al. 2014).

**Figure 1:**
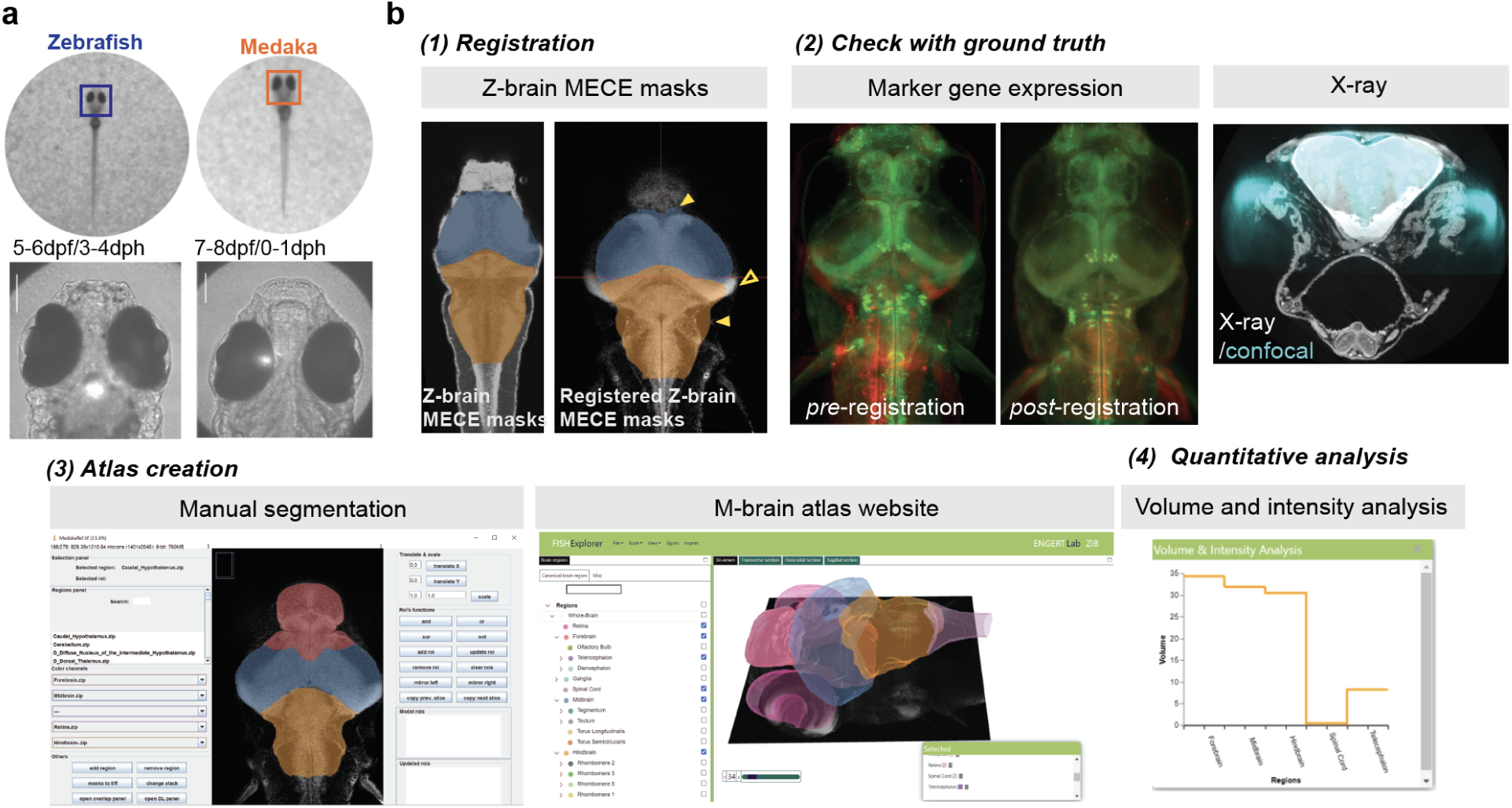
Workflow of creating an atlas for larval medaka by using the interactive atlas-based platform. (a) Comparison of whole bodies and heads of teleost larvae, zebrafish (5 dpf/3 dph) and medaka (8 dpf /1 dph). Scale bars: 150um. (b) Workflow for creating the M-brain atlas to the interactive atlas platform: (1) The MECE masks of the Z-brain were registered to the medaka reference brain. Yellow, filled arrowheads indicate over-registration, while the hollow arrowhead indicates under-registration. (2) By visualizing marker gene expression (left) and X-ray images (right), the ground truth was confirmed. (3) A custom-developed Fiji plug-in, Neuro Mask Organizer (NeMO), allowed us to define and segment medaka MECE masks automatically using a 2D-UNet (left). Then the M-brain atlas was implemented (right). (4) The interactive atlas includes the function for visualization of the volumes and intensities of brain region masks. Here, the volumetric comparison of large MECE regions in medaka is shown (orange).

To establish the M-brain atlas, we extended the existing FishExplorer atlas platform that contains the Z-brain atlas. First we defined anatomical masks for medaka larvae by adhering to the following steps: (0) creation of a reference brain for medaka larvae by using four fish individuals; (1) registration of the Z-brain MECE masks to the medaka reference brain; (2) determination of the ground truth by neurotransmitter expression; and (3) final editing of the masks using a new, customized Fiji plug-in, Neuro Mask Organizer (NeMO), for segmenting brain regions (see Methods) (Figure 1b).

For generating the reference brain stacks, we incorporated Hoechst dye staining, immunostaining such as total-ERK (Randlett et al. 2015), and HuC promoter labeling (Ahrens et al. 2012) and integrated them in our multi-channel reference brain (Supplementary figure 1), allowing us to support a large range of analyses. For registration, we used the Advanced Normalization Tools (ANTs) (Avants et al. 2014) which offers algorithms for both rigid and non-rigid alignment of image stacks.

Next, the zebrafish MECE masks from the Z-brain atlas were registered onto the medaka reference brain to generate preliminary medaka MECE masks (Figure 1b). Although the overall quality of the registration of the large region masks was good, over- and under-registration of the masks can be observed at boundaries (Figure 1b). Regarding the small brain regions, most of them were not registered at the precise location. To correct this, we followed a semi-automated technique that combined manual labeling with a machine learning approach. Since manual labeling of the masks is tedious and requires a lot of time from expert users, we added a machine learning algorithm where we trained a 2D-UNet network (Ronneberger et al. 2015) that is based on a deep learning model for the segmentation of the big regions (Supplementary Figure 2). After the initial segmentation of the masks by the experts, we applied a new segmentation tool, Neuro Mask Organizer (NeMO), which was developed as a custom Fiji plug-in (Figure 1b, Supplementary Figure 2). The plug-in includes several utility functions, e.g. comparing masks with registered regions, searching for overlaps between regions, and copying masks into continuous slices. For each registered mask derived from the Z-brain atlas, initially a few slices with large intervals were labeled manually using our plug-in. Following this, machine learning models were trained on those slices, and masks were generated for the unprocessed slices using the trained model. We found that 20-30% manual labeling of regions distributed across the whole stacks was enough to complete segmentation of the whole brain (Supplementary Figure 3). In total, we segmented 21 relatively large regions using deep learning (see Methods section for detailed description of the methodology). In the end, the deep learning-based predicted masks were fused and final region masks were generated using the new plug-in. In order to fix the displacements and/or sizes of the small anatomical regions, we validated their location and size with the marker-genes expression patterns (see Appendix, Supplementary Figure 4). Finally, an X-ray dataset was generated and used to verify the locations of all the neuromasts – a specialized set of sensory organs that most teleost fish use to sense small perturbations in water currents (Figure 1b and Supplementary Figure 5).

### Establishment and comparison of MECE region masks

Using the interactive brain atlas platform FishExplorer, hosting both the zebrafish and the medaka larvae brain atlases, we first performed volumetric comparison of MECE regions between zebrafish and medaka. To this end, we assigned a unique identifier for each MECE region and listed its volume (**Figure 2, Table 1**). Next, we normalized the mask volumes by the total brain volume (except for the retina) of each fish species and quantified relative volumes. Using this approach, we found that the midbrain in medaka is bigger than in zebrafish, whereas the hindbrain is bigger in zebrafish than in medaka (**Figure 2b**). In the smaller subsets of MECE masks, we confirmed that 113 of the MECE masks exist in both zebrafish and medaka. While we found medaka-unique neuromasts in the midbrain (neuromast SO4), a few of the reticulospinal cord neural clusters (MiR1 and MiR2) are missing in medaka (**Figure 2**, see the **Appendix** for more details).

**Figure 2:**
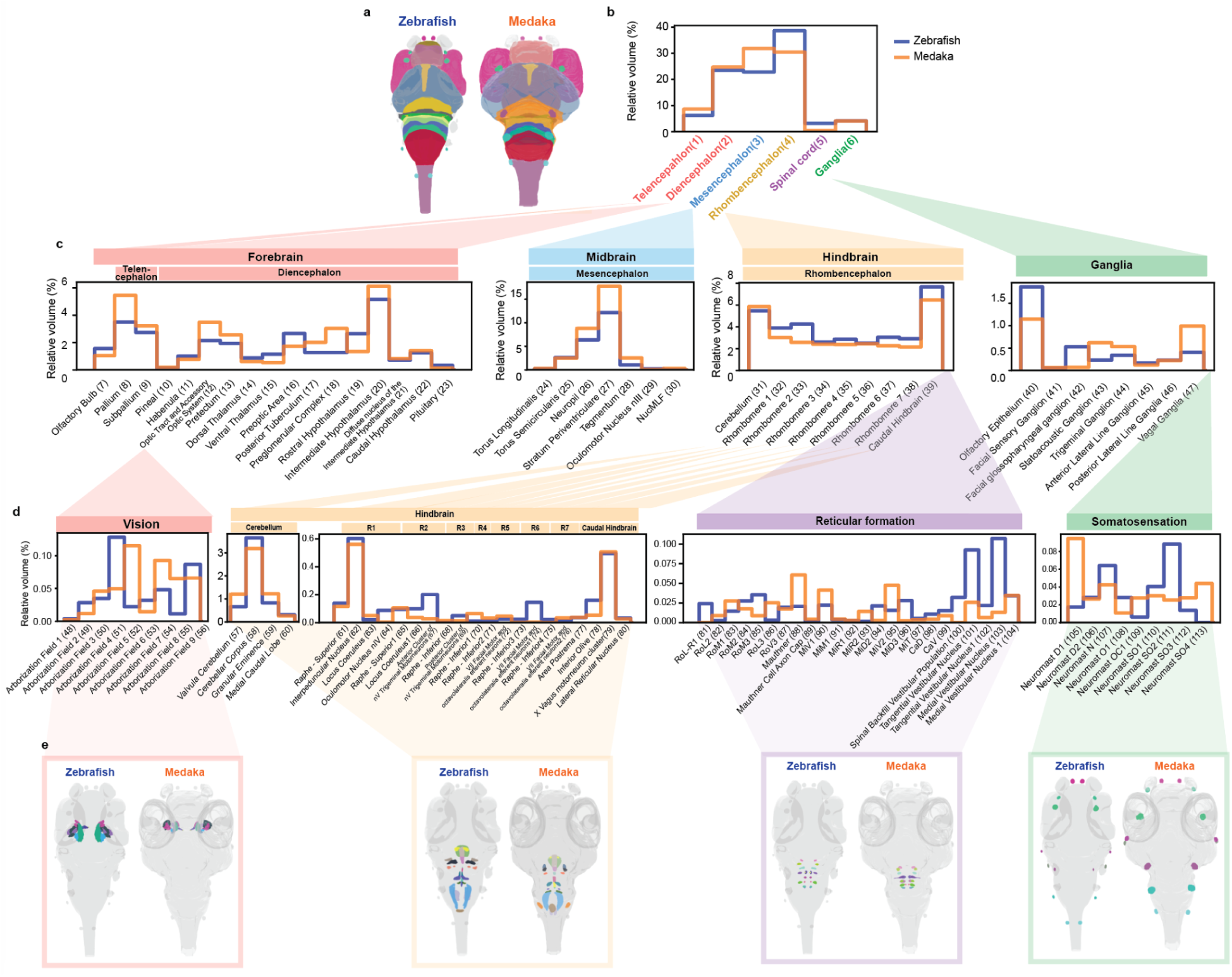
Volumetric comparison of MECE region masks of teleost larvae of zebrafish and medaka. (a) 3D representation of MECE region masks of zebrafish (left) and medaka larvae (right). The colors of the masks correspond to the regions in (**b**). (**b**) Red: forebrain (Telencephalon, Diencephalon), blue: midbrain (Mesencephalon), orange: hindbrain (Rhombencephalon), purple: spinal cord, green: ganglia. Numbers after the names of MECE masks stand for the assigned IDs, which is assigned from the dorsal to ventral, and anterior to posterior directions. (**c, d**) Volumetric comparison of MECE subregion masks. Manhattan plots show relative volume across MECE masks in zebrafish (blue) and medaka (orange). Numbers after the name of MECE masks represent the MECE mask IDs. (**d**) Nested small MECE masks are shown. (**e**) 3D representation of MECE masks in (**d**). Abbreviations - see the legend of Supplementary figure 5. Reticular formation: M (Mauthner cell), RoM (rostral of Mauthner cells) clusters, MiM (middle cells in middle area), MiV (ventral cells in middle area), MiD (dorsal cells in middle area), MiT (T-shaped axon in middle area), MiR (rostral cells in middle area), MeM (projection in mesencephalon, medial), “MeLm (projection in mesencephalon, lateral medial), MeLr (projection in mesencephalon, rostral), MeLc (projection in mesencephalon, caudal), RoV (projection in rhombencephalon, ventral), RoL (projection in rhombencephalon, lateral), CaD (dorsal cells in caudal area), CaV (ventral cells in caudal area), NucMLF (nucleus of medial longitudinal fasciculus), V (vestibular clusters). Neuromast N (Nasal neuromasts), SO (supraorbital neuromasts), O (optic neuromasts), OC (occipital neuromasts), MI (middle neuromasts) and D (dorsal neuromasts).

### Establishment and comparison of labeled neural clusters

The expression patterns of neurotransmitters are also useful to identify the homology of brain regions across vertebrates, in addition to assigning potential functional properties. For a quantitative comparison between zebrafish and medaka, we selected the following major neurotransmitters and neuropeptides (**Figure 3**): GABA, glutamate, serotonin (Lillesaar 2011), dopamine (Matsui 2017), noradrenaline (Tay et al. 2011), and oxytocin (Wee et al. 2019).

**Figure 3:**
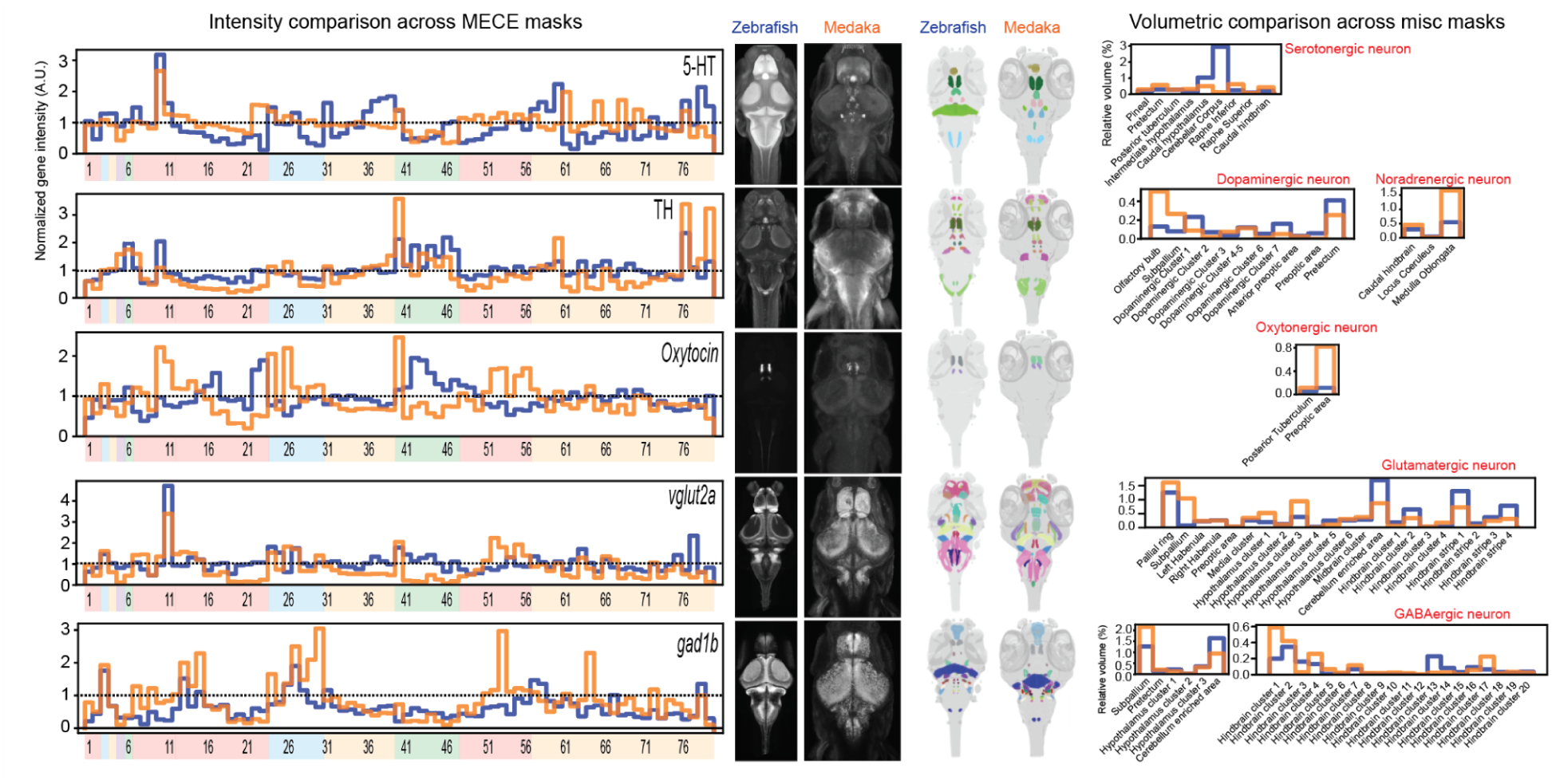
Comparison of selectively labeled subclasses of neuronal populations. Expression level of the basic neurotransmitters and neuropeptides. (Left) Intensity plot across the whole brain within the MECE masks. The numbers and color code on the horizontal axis correspond to the identification numbers of the MECE masks in Figure 2. (Middle left) Maximum intensity plot of staining stacks of zebrafish and medaka. (Middle right) 3D representation of gene clusters with different colors indicating different clusters. (Right) Volumetric comparison of gene clusters (blue: zebrafish, orange: medaka). 5-HT: 5-hydroxytryptamine, TH: tyrosine hydroxylase, vglut2a: vesicular glutamate transporter 2a, gad1b: glutamate decarboxylase 1b (see **Appendix** for details).

First, signals visualized by various methods (immunostaining, *in situ* hybridization, and transgenic lines) were quantified across the whole brain, using MECE masks (**Figure 3 left, middle**). We found that many of the expression patterns of marker genes are similar between zebrafish and medaka. In a few cases, where the expression is limited to a small subpopulation of a large MECE region masks (e.g. sparse Oxytocin clusters in the posterior tuberculum and preoptic area), the expression value of MECE region mask got decreased by surrounding non-expressing cells. In order to identify specific neural subpopulations that express marker genes, MISC masks were generated in medaka, as performed for Z-brain atlas in a previous paper (Randlett et al. 2015). Transforming labels into MISC masks required expert knowledge (see ***Supplemental information***). We then compared the relative volumes of the MISC masks between zebrafish and medaka (**Figure 3, right**), and found most of the clusters to be homologous, which suggests strong conservation of gene expression across species. However, we found the size of some clusters to be different, suggesting potentially different connectivity and function of the neural circuits formed by these clusters.

### Comparison of the telencephalon in teleost larvae

The interactive multi-species brain atlas platform allowed us to analyze teleost larval brains and address questions in evolutionary diversity. One of the brain areas with the highest diversity across vertebrates is the telencephalon. The mammalian telencephalon has been intensely studied; it includes the amygdala, the hippocampus, as well as the neocortex, and the precise function and homology of these regions in other amniotes is still largely unknown (Hain et al. 2022). Also previous studies suggest a big diversity of anatomical regions inside the telencephalon even across teleost species, and many studies have been done in the adult teleost brain (Mueller and Wullimann 2009); Isoe et al 2023, (Tibi et al. 2023). However, comparative analysis in larvae still remains largely elusive.

To illustrate and characterize the diversity of the telencephalon in larval zebrafish and medaka, we labeled and generated new MISC (miscellaneous) masks based on the density of synapses by leveraging a presynaptic marker, synapsin, which is ubiquitously expressed in all vertebrates. Anti-synapsin immunostaining suggests different compartment patterns of computation in the telencephalon between two species (**Figure 4a,b**). By quantifying the expression, we found medaka to display stronger synapsin-positive subregions in the dorsal area of the pallium (Dd), as well as in the medial area of the pallium (Dm4). On the other hand, no strong synapsin staining was detected in the medial part of the pallium (Dm) in zebrafish, but the posterior area of the pallium (Dp) showed strong staining (**Figure 4c**). Next, to detect gene profiles in the subregions of the telencephalon, marker gene expression was quantified in the pallium using the newly generated MISC masks (**Figure 4c**). We found that the expression of dopaminergic signals is similar between zebrafish and medaka, with the exception of the Dp region. Further, the serotonergic signals are highly expressed in Dd in medaka, whereas they are most detectable in Dp in zebrafish. Glutamatergic expression is higher in most regions in medaka, and the GABAergic expression pattern is similar in both species.

**Figure 4:**
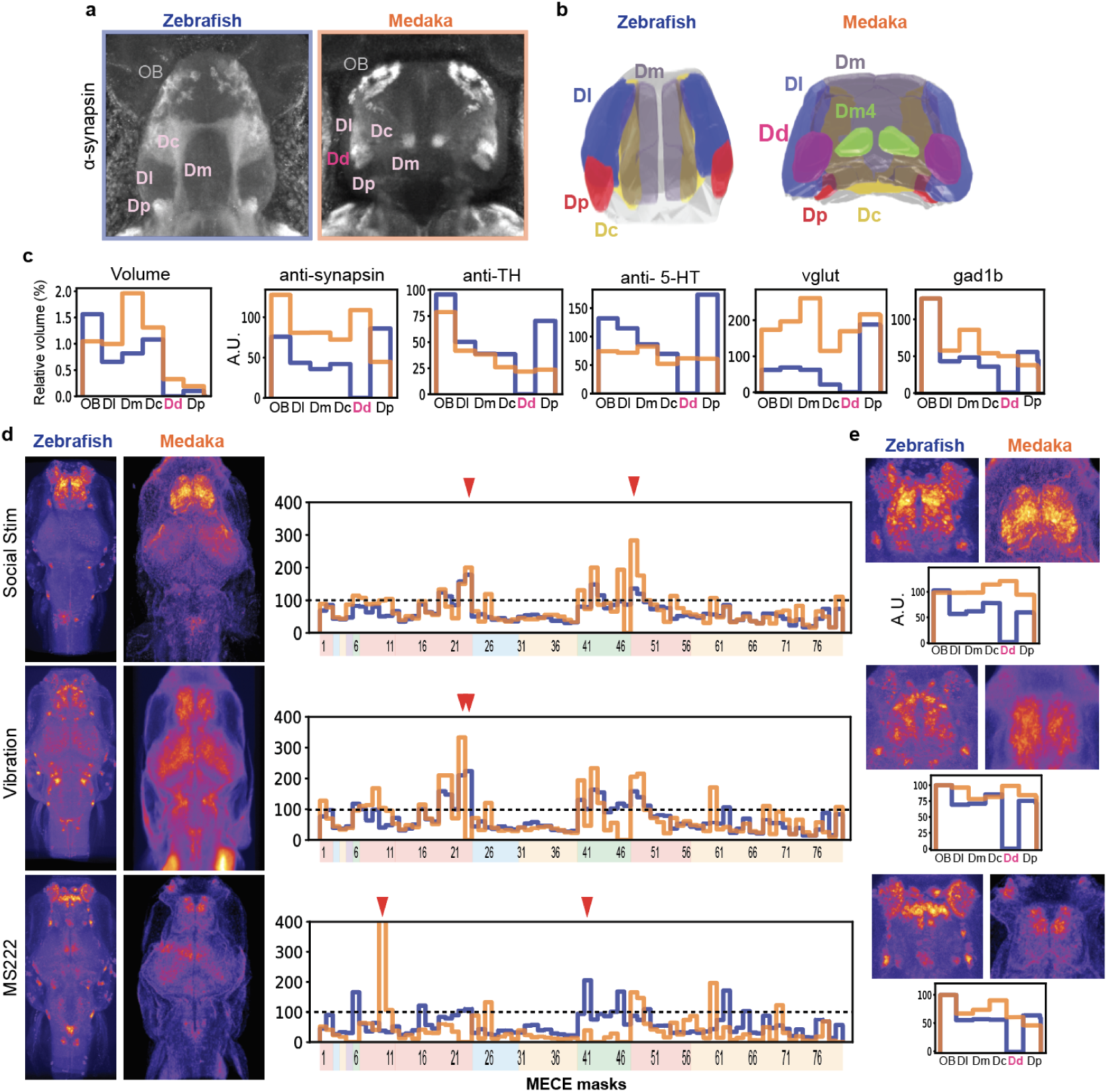
Comparative analysis of the telencephalon and activated neural populations. (**a**) Comparison of the telencephalon. Anti-synapsin immunostaining exhibits anatomical subregions in the telencephalon. In zebrafish, a high density of synapsin is detected in Dc and Dp, while in medaka a high density of synapsin is observed in Dd and a subregion in Dm (Dm4 is a posterior subregion in Dm whose synapsin expression is intense). OB: the olfactory bulb, Dc: the central part of the pallium, Dd: the dorsal part of the pallium, Dl: the lateral part of the pallium, Dm: the medial part of the pallium, Dp: the posterior part of the pallium. (**b**) MISC masks were generated to study the telencephalon based on the anti-synapsin and Hoechst staining. (**c**) Manhattan plots comparing the volumes of MISC masks and neurotransmitter expressions. A.U.: arbitrary unit. (**d**) Visualization of the neural activity after three conditions with *cfos* staining by *in situ* hybridization. The maximum projection of optical sections in the horizontal direction is shown (left). Comparisons of *cfos* intensities across MECE regions are shown with a Manhattan plot (zebrafish: blue, medaka: orange). The numbers and color code written on the horizontal axis correspond to the identification number of MECE masks in Figure 2. Red triangles indicate the highest *cfos* intensity in zebrafish and medaka. (**e**) Magnification of the *cfos* staining in the telencephalon (top); comparisons of *cfos* intensities across MISC masks in the telencephalon (bottom; zebrafish: blue, medaka: orange).

### Comparison of neurally activated regions by c-fos staining

To leverage the MECE and MISC masks in a behavioral neuroscience context, we compared neural activity when animals were stimulated through different sensory modalities. Specifically, we selected full anesthesia by immersion in MS222 as a control, vibrational tactile stimulation and exposure to a group of conspecifics as a social stimulus. After these respective treatments, we visualized the activated cells by detecting the expression of the immediate early gene, *cfos*, by *in situ* hybridization (**Figure 4d**). Remarkably, we found the expression of *cfos* in all of the modalities to be quite similar in zebrafish and medaka. MS222 immersion resulted in the silencing of most brain regions, with the exception of the neuromasts, the olfactory bulb (OB), the area postrema and vGlut cluster 1 in the mesencephalon in both species. Vibrational stimuli activated the subpallium and glutamatergic clusters in the hindbrain in both zebrafish and medaka, whereas social stimuli were found to elicit strong signals in the olfactory epithelium and the neuromasts in zebrafish, while stronger signals were observed in the optic tectum in medaka larvae. In addition, very strong signals in the telencephalon, especially in the pallium, were observed in both species after social stimulation.

Next, we focused on the telencephalon and compared the neural activity in its subregions by using the MISC masks generated in both species (**Figure 4e**). We found that *cfos* expression was pretty similar between zebrafish and medaka, with MS222 immersion leading to strong activation of the medial OB, whereas vibrational stimuli elicited only weak signals compared to those detected in other brain regions. However, within the telencephalon we found that OB showed strong signals in both species, and Dd in medaka. Social stimulation elicited very strong signals in the telencephalon of both species, with the Dd region dominating in medaka, whereas subpallial signals were stronger in zebrafish.

### Data integration to the Multi-species platform

The finalized medaka data-set was added to our interactive web-based FishExplorer platform as an integrated feature (M-brain atlas, https://fishexplorer.zib.de/medaka/lm). On this combined website, the reference brain stacks are available for users to register their own dataset (**Figure 5**, code for registration can be found in the tutorial). After registration, users can upload their registered stacks to their own private account of the website and visualize them with MECE and MISC masks in 2D and 3D. If necessary, users can draw new MISC masks with the new Fiji plug-in, NeMO, and upload them in their private account. After publishing the stacks and/or masks, the user can share the uploaded stacks with the community and they will be transferred from the private account to the public database on the FishExplorer website. The interactive nature and easy scalability of the atlas facilitates the engagement of the teleost community and encourages it to contribute to its continuous expansion.

**Figure 5.**
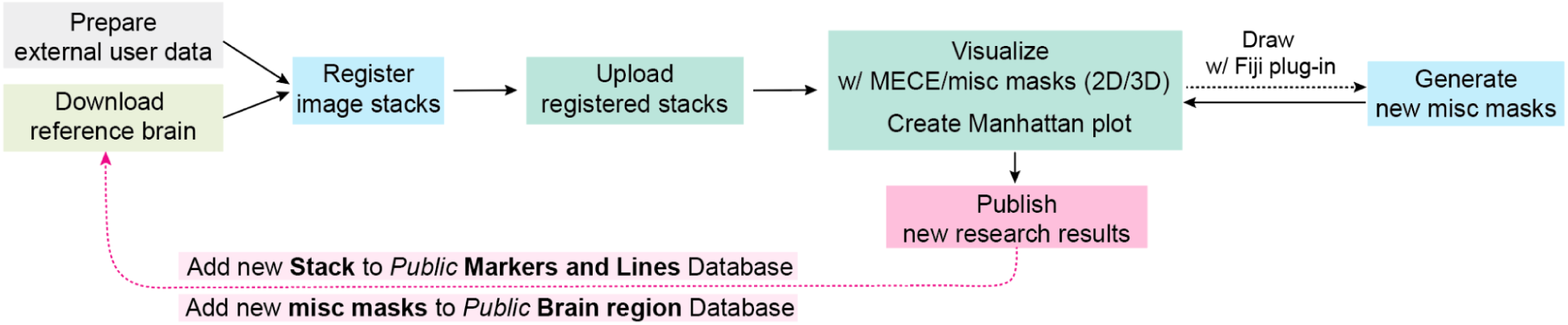
Flow chart for data integration to the atlas platform. Workflow for users to analyze their own data. Users can download the reference brain stacks and register their dataset to the reference stacks with Python scripts. Then, the registered brain stacks can be uploaded to the website of their private account and users can visualize them with existing MECE and MISC masks. Users can also draw and generate new MISC masks with the Fiji plug-in, NeMO, and upload these masks to their private account. When a paper is published with the user’s dataset, the new stacks are added to the public markers and lines database, and new MISC masks are added to the public brain region database. The blue boxes in the chart indicate activities that users perform in their own workspace, while the green boxes indicate the activities that users can perform in the atlas platform.

## Discussion

Here we presented a new multi-species brain atlas platform for larval teleost that extends the existing FishExplorer platform containing a zebrafish atlas by adding the necessary functionality for cross-species comparison with medaka fish, a phenotypically similar but evolutionary distant teleost species. To facilitate engagement of the teleost science community, we developed a custom-build Fiji plug-in, NeMO, providing efficient utility and deep learning functions to interactively generate brain region masks for integration into the atlas platform.

This platform allows to quantify relative volumes, as well as gene expression profiles across the whole brain. The relative volumetric comparison across MECE regions allowed us to speculate on evolutionary mechanisms that might have led to brain specifications depending on the natural environments where the species have evolved. We hypothesized that the more sensory modalities are used, the larger the brain regions corresponding to the modality have evolved to be, and vice versa. For example, some teleost species exhibit volumetric diversity in brain regions related to the sensory modality the species utilize. As such, weakly electric fish have evolved a very large cerebellum that is used for predictive electric discharge calculations in hunting and communication (Sukhum, Shen, and Carlson 2018). Mexican cavefish (*Astyanax mexicanus*) are blind and have a relatively small optic tectum compared to surface *A. mexicanus* since they have evolved in a lightless environment for ∼20.000 years (Jaggard et al. 2020). Additionally, they also have a bigger hypothalamus and telencephalon (Loomis et al. 2019), possibly since non-visual navigation is more computationally intensive. Here, we found that medaka has a larger midbrain, with an especially enlarged stratum periventricular in the optic tectum, and a smaller hindbrain than zebrafish. This might be because medaka naturally inhabit a clear water environment which potentially leads to a higher computational load when processing visual information. In contrast, the hindbrain in zebrafish might have evolved to a larger size, because they possibly adapted to the need to respond quickly and with more complex motor patterns to relevant visual stimuli that appear in close proximity in the murky waters in which they evolved.

Additionally, we found a pair of neuromasts that exist in medaka but not in zebrafish. This phenotype has also been observed in various cavefish species, yet is difficult to link to any adaptive behavioral relevance (B. Chen et al. 2022). The observation that neuromasts get activated in many treatments in zebrafish, while we did not observe strong neural activity in neuromasts in medaka, could be related to a general propensity for non-visual navigational strategies adapted to the murky environment of the Ganges delta, the evolutionary home of the zebrafish.

Our interactive brain atlas platform provides tools for users to generate new MISC region masks, which allows for an unrestricted specification of any particular region of interest. Here we have created masks for the telencephalon, which has recently received particular attention in gene profiling and function analysis. A recent paper showed that there is an epigenetically distinct region in the dorsal pallium (Dd) in medaka larvae (Isoe et al., n.d.), which had not been confirmed to exist also in zebrafish larvae, although juvenile and adult zebrafish have distinct functional compartments in the pallium (Bartoszek et al. 2021). Here, we compared the gene expression and neural activity in different contexts in the compartment of the telencephalon in larval medaka and zebrafish, and we found that the Dd region has a unique expression pattern of marker genes.

Using *cfos hybrid chain reaction (*HCR) (Choi et al. 2018) staining, we found that the brain regions activated by different stimuli are mostly the same in zebrafish and medaka, which is interesting given the large evolutionary distance between the two species. Consistent differences in medaka and zebrafish were only observed in the olfactory epithelium and the neuromasts. Under all conditions, *cfos* was highly expressed in the neuromasts and the olfactory epithelium in zebrafish, but not as strongly in medaka. The olfactory epithelium was also found to be larger in zebrafish, suggesting that zebrafish have adapted to a more murky environment by relying more on olfactory information than medaka. We found that the area postrema is activated by MS222 in both species, which is consistent with previous findings that this area is activated by chemical sensation (Wee et al. 2019; Tsukamoto and Adachi 1994). The vibrational assay confirmed a conserved activation of hindbrain glutamatergic neurons in both species. The social stimuli uncovered a conserved role of the telencephalon in both species. Further analysis of regions within the telencephalon suggested that the Dd regions in medaka are differentially activated compared to other pallial regions, suggesting a possible functional significance of this region. Cross-species analysis of more complex behaviors will require a detailed description of the adult and later developmental stages in both species, necessitating an extension of our atlas platform to developmental stages at older ages.

Even though this atlas platform worked well in medaka and zebrafish, they are phylogenetically very distant (250 Mya), we estimate that it would be hard to apply for species in different animal classes. The reason it worked well in medaka and zebrafish is that the basic brain structure is mostly conserved in the same animal classes. Also if there is an evolutionary completely different brain region, it would be difficult to identify it by morphing the masks by itself, unless the researchers perform stainings of several marker genes, as we performed in the current paper. We expect the researchers will use our manual annotation function in our Fiji plug-in, NeMO, to generate a new mask for the unique brain region. NeMo contains the mask generating function utilizing machine learning. This function works well in the big brain regions, but it is not optimal for the small regions, which will be supported by the manual annotation function, too.

The study of evolutionary diversity and principles of behavioral mechanisms using comparative approaches in neuroscience and neuroethology has recently attracted more attention with the advances in imaging technologies, genetics and genomics. Our interactive, multi-species, community atlas has the capability to accommodate, with moderate effort, datasets collected by research groups that study zebrafish and medaka neuroscience across the world. At the next stage we plan to extend our platform to include other teleosts, such as danionella, and add other developmental stages to facilitate the study of developmental aspects in addition to interspecies comparisons.

## Materials and Methods

### Animal experiments and ethics

All experiments followed institution IACUC protocols as determined by the Harvard University Faculty of Arts and Sciences standing committee on the use of animals in research and teaching. The animal experimentation protocols, 25-03 and 22-04 were submitted and approved by this institution’s animal care and use committee (IACUC). This institution has Animal Welfare Assurances on file with the Office for Laboratory Animal Welfare (OLAW), D16-00358 (A3593-01).

We followed The ARRIVE guideline 2.0. Sample size is described in the method section. Inclusion and exclusion criteria: when the registered imaging stacks didn’t align well with the reference brain, we did not include those stacks for further analysis. Randomization: for the analysis of *cfos* staining experiments, fish samples were imaged in random order with the same imaging parameters. Blinding/masking, outcome measures: For quantitative analysis of the gene expression, we masked the label of genes until the result interpretation.

### Fish maintenance

All medaka fish and zebrafish wild type and transgenic lines used in this research were raised in filtered fish facility water on a 14 h light, 10 h dark cycle at 28°C. Under 28°C incubation, medaka hatch at 6-7 days post-fertilization (dpf), while zebrafish hatch at 2 dpf (Signore et al. 2009). We used 0 or 1 days post-hatch larvae for medaka and 5 -7 dpf larvae for zebrafish.

### Transgenic lines

Tg(vglut:GFP) line was generated in the Higashijima lab and provided by NIBB. Tg(Ath5:GFP) line was generated by the Wittbrodt lab (Del Bene et al. 2007). Tg(HuC:loxp-DsRed-loxp-GFP) in ST3 mutant background was made by crossing Tg(HuC:loxp-DsRed-loxp-GFP) (Okuyama et al. 2013) with ST3 mutants (Wakamatsu et al. 2001). Tg (ST3-eyesBlack) was established by crossing ST3 mutants and drR, then embryos were screened when they don’t have autofluorescence around the head but have pigment cells in the eyes.

### Immunochemistry and in situ hybridization

We followed the immunostaining procedure from the previous study (Randlett et al. 2015). Information on primary antibodies can be found in **Supplementary Table 1**. Secondary antibodies conjugated with Alexa fluorophores (Life Technologies) were diluted 1:250. Hoechst dye was diluted 1:1000.

*In situ* hybridization experiments were performed being followed by the procedures previously published (Vauti et al. 2020; Shainer et al., n.d.). In brief, after fixing fish larvae in 4% PFA/PBS overnight, larvae were dehydrated with 100% methanol. Then we washed with ethanol and performed permeabilization with ethanol/xylol solution. Then we rehydrated with a sequence of ethanol/H2O with 0.1% Tween-20. Then we permeabilized larvae with 80% acetone/H2O at -20C for 30 min and washed larvae with PBST (0.1% Tween-20, PBS). We pre-hybridized the larvae with a prewarmed hybridization buffer for 4 hours and incubated them with HCR probe for 60 hours at 37 C. Then larvae were washed with a 5 x SCCT buffer and incubated with an amplification buffer for 30min. After that, larvae were mixed with hairpin that heated up at 95 C for 90 seconds, and incubated overnight at RT. After washing with 5 x SCCT, we cleared the larvae with a series of Glycerol and stored at -20C. We always stained with *snap25,* which was used as a whole-neuron marker for the registration. We ordered HCR-RNA bundles from Molecular Instruments.

Quantitative comparison of gene expression between species allows us to assess differences in the function of neural circuits and brain regions. The difficulty in comparing signal intensity was to normalize and compare the signals across regions located in different depths and between different species. Since slices close to the microscopy objective tend to be detected with higher signals than slices far away, we normalized the gene intensity values by dividing them by signals of pan-cell or pan-neuronal markers, such as anti-tERK immunostaining or *snap25 in situ* hybridization (HCR). To compare signals between species, we normalized with the averaged intensity of the entire stacks.

**Supplementary Table 1.**
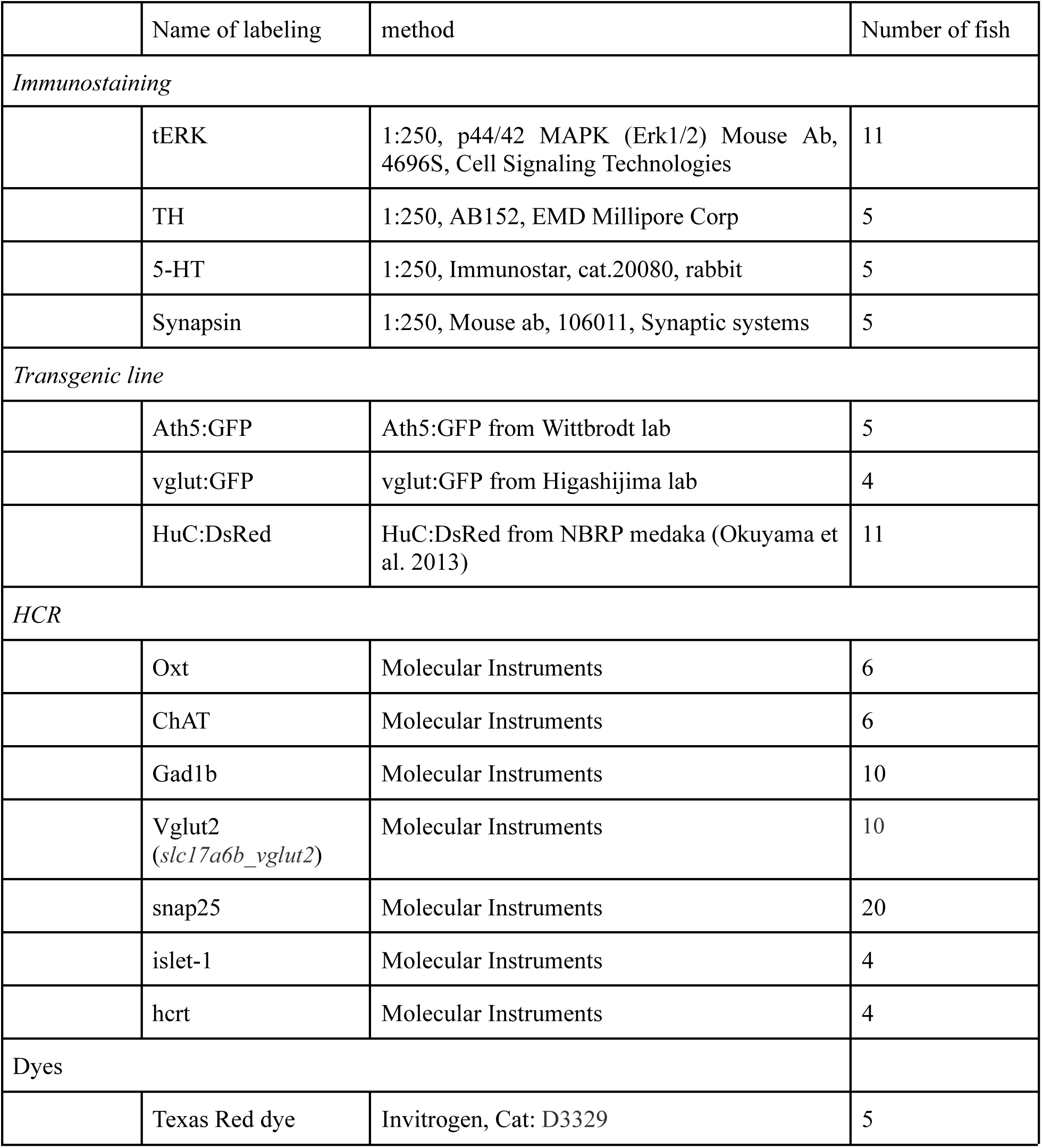
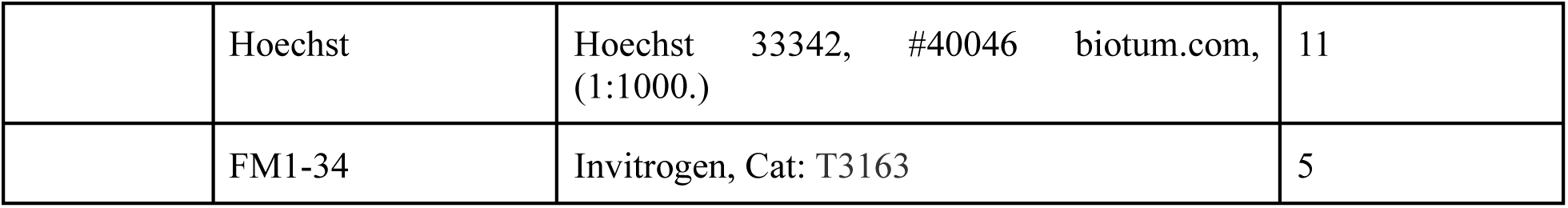
Staining protocol.

### Backfill labeling

We first embedded larvae into 2% agarose gel sideways down. After scooping out the gel above the fish, Texas Red crystals were injected into the floor plate of larvae with a tungsten needle. We removed the larvae from the gels immediately after injection and placed them in the fish water. The next day, we fixed the larvae in 4% PFA with Hoechst (1:1000) overnight. After washing with PBS-Tx, we embedded the fish larvae in agarose (2% in PBS) and performed imaging under a confocal microscope (Zeiss LSM780).

We identified the Mauthner cells by their shape, and based on the position relative to the Mauthner cells, which are located in rhombomere 4, we identified other structures, such as nucleus of medial longitudinal fasciculus (NucMLF), RoM cells and MiV cells (Metcalfe, Mendelson, and Kimmel 1986).

### FM1-34 staining

We transferred fish larvae into FM1-34 (3uM) solution for 30 sec. Then fish were transferred to fish water for 10 min and washed several times. We fixed the larvae in 4% PFA with Hoechst dye (1:1000) overnight. After washing with PBS-Tx, we embedded the fish larvae in agarose (2% in PBS), and we performed imaging under a confocal microscope (Zeiss LSM780).

### X-ray imaging

The sample was stained to enable both electron microscopy and X-ray imaging following a protocol modified from (Tapia et al., 2012). Unless noted, all steps were performed at room temperature (RT). At one day post hatch the larva was anesthetized and embedded in agarose, we replaced the water by a dissection solution (64 mM NaCl, 2.9 mM KCl, 10 mM HEPES, 10 mM glucose, 164 mM sucrose, 1.2 mM MgCl2, 2.1 mM CaCl2, pH 7.5) supplemented with 0.02% tricaine (Hildebrand et al., 2017). We then cut small slits in the agarose to expose the eyes, performed bi-lateral enucleations and tail resection to enhance ultrastructural preservation and heavy metal staining. We used a custom-made hook that was carefully inserted behind the eyes to prevent brain damage. The larva was immediately transferred to a fixation solution at 4°C (2.5% glutaraldehyde, 0.1M cacodylate buffer supplemented with 4.0% mannitol at pH7.4). Cacodylate buffer: 0.3M sodium cacodylate, 6mM CaCl2, pH7.4. To improve the fixation the tissue was rapidly microwaved (cat. no. 36700, Ted Pella, equipped with power controller, steady-temperature water recirculator and cold spot) in the fixative solution (this step lasted <5 min after initial transfer into fixative). The microwaving sequence was performed as in (Tapia et al., 2012): at power level 1 (100 W) for 1 min on, 1 min off, 1 min on then to power level 3 (300 W) and fixed for 20 s on, 20 s off, 20 s on, three times in a row. Fixation was then continued overnight at 4°C in the same solution. The following day, the sample was then washed again in 0.5x cacodylate buffer (3 exchanges, 30 min each before osmication (2% OsO4 in 0.5x cacodylate buffer, 60 min). After a quick wash (< 1 min) in 0.5x cacodylate buffer the sample was reduced in 2.5% potassium ferrocyanide in 0.5x cacodylate buffer for 60 min then washed with filtered H2O (3 exchanges, 30 min each) and then incubated with 1% (w/v) thiocarbohydrazide (TCH) in filtered H2O (and filtered with a 0.22um syringe filter before use) for 20 min to enhance staining (Hua et al., 2015). Due to poor dissolution of TCH in water, the solution was heated at 60°C for ∼90 min with occasional shaking before filtering and then placed at RT for 5min before the incubation step. The sample was then washed with filtered H2O (3 exchanges, 30 min each) before the second osmication (2% OsO4 in filtered H2O, 60 min) and then washed again (3 exchanges, 30 min each). Then, en-bloc staining was performed overnight using 1% uranyl acetate in filtered water at 4 °C. The solution was sonicated for 90 min and filtered with a 0.22um syringe filter before use. Steps involving uranyl acetate were performed in the dark. The following day, samples were then washed with filtered H2O (3 exchanges, 30 min each). Next, the samples were dehydrated in serial dilutions of ethanol (25%, 50%, 75%, 90%, 100%, 100% for 10min each step) then in propylene oxide (PO) (100%, 100%, 30min each step). Infiltration was performed using LX112 epoxy resin with BDMA (21212, Ladd) in serial PO dilutions steps, each lasting 4h (25% resin/75% PO, 50% resin/50% PO, 75% resin/25% PO, 100% resin, 100% resin). Samples were mounted in fresh resin in a mouse brain support tissue (Hildebrand et al., 2017) with the head exposed to facilitate cutting by avoiding the sample to sink to the bottom of the resin molds. Mouse tissue was fixed using standard procedures (Fang et al., 2018) then cut into 2-3mm wide cubes which were pierced using a puncher (0.75mm, 57395, EMS) to insert the larva. The cubes were stained along with fish samples using the protocol described above except that the uranyl acetate overnight step was performed at RT. The samples were then cured with support tissue during 3 days at 60°C. For all steps, a rotator was used. Aqueous solutions were prepared with water passed through a purification system (Arium 611 VF, Sartorius Stedim Biotech). The protocol lasted 5 consecutive days including surgery, fixation, staining, and resin embedding followed by 3 days of resin curing. The block was finally trimmed to expose the head of the fish for X-ray imaging (Xradia 520 Versa, ZEISS). Imaging was performed at 40 kV, 76 uA, 20X optical magnification, 0.35 um isometric voxel resolution and lasted 96 hours (100 seconds per image).

To register the brain masks using X-ray imaging dataset, we performed the annotation using Fiji plug-in, BigWarp. Since X-ray imaging dataset cover larger areas of the fish, such as gills and other organs, we registered the masks from the reference brain onto X-ray data, but not the other way around.

### Segmentation of brain region masks with Fiji plug-in, Neuro Mask Organizer (NeMO)

For the segmentation of the brain regions, we implemented a tool that is integrated as a plug-in in the ImageJ/Fiji platform (Reuden et al., 2017). The main motivation behind using this custom plug-in compared to the state-of-the-art segmentation editors available in Amira, VAST (Berger et al., 2018) etc. is to create various utility functions. These include copying masks to the left/right brain hemisphere, comparing masks with registered zebrafish masks, and finding overlaps between regions, to enable fast and interactive ground-truth labeling. The new masks/regions of interests (ROIs) were labeled using polygon and free hand selection tools of ImageJ. The plug-in (**Supplementary Figure 2**) consists of several panels: (1) Region panel displays a list of brain regions (indicated by a red arrow head).Users can select/deselect a brain region by clicking on an item in the list. The panel also shows the name of the currently selected region and the region of interest in a plane. (2) Color channel panel (indicated by yellow arrow head). In order to label a selected region, users often need boundaries of adjacent regions. This is accomplished by loading adjacent regions as a colored channel. (3) Other panels (indicated by purple arrow head). This allows users to open two important sub-panels: the overlap panel, which provides a list of overlaps between the selected region and the loaded regions in the colored channel panel. It also provides helper functions to fix the overlaps and deep-learning panel that allows users to create the training dataset and fuse the model predictions. Users can choose different options for creating a training dataset, e.g. training on whole images or image sections, percentage of labeled data, split strategy etc. (4) ROI panel (indicated by blue arrow head). It consists of two sublists, namely labeled ROIs for a selected region in the current slice and predicted ROIs in the current slice. (5) ROI functions (indicated by green arrow head). They enable users to perform boolean operations between a selected ROI and the currently labeled ROI. It also enables users to copy a selected ROI to the other hemisphere or the next/prev slice. The plug-in will be available on github including detailed instructions. We defined MECE masks and MISC masks based on the cell localization by Hoechst staining and gene expressions visualized by immunostaining, in situ hybridization and transgenic lines. Please see the **Appendix** for more details.

### Segmentation of brain regions with deep learning

Manual segmentation of brain regions requires a lot of time and expert effort. Our aim here is to reduce the effort of manual segmentation by using deep learning models. For this purpose, we used the following dataset and methodology.

We recruited the Hoechst dye-stained reference brain to perform the segmentation experiments. The original dimension of the reference stack was 1401x2045x278 voxels. We upsampled the stacks to 2045x2045x278 voxels to obtain an isotropic voxel size. For performing our experiment, we downsampled the stack to 512x512x278 voxels to increase the receptive field of the network. We applied the segmentation to 21 large brain regions. To test our method, the expert manually performed complete labeling of 8 regions (**Supplementary Figure 1a**) using our Fiji plug-in, NeMO. The remaining 13 regions initially remained unlabeled and were later labeled based on experimental results of these 8 regions.

First we checked how much manual labeling is sufficient to train the network model. From the 8 fully labeled binary-segmented brain regions, we generated training data with 20%, 25%, 30% and 35% of them using the deep learning panel of the Fiji plug-in, NeMO. We did not keep any data for validation because that would have required a further 10-15% of segmented data. The major drawback of not employing validation data would be an overfitted model but our goal is to reduce the manual efforts rather than to prevent overfitting. To increase the set of training data, we applied data augmentation. In particular, we applied random shifts with range of [-50, 50] pixels, [-5, 5] degree rotation, random flips, shear of 0.06 degree in counter clockwise direction, zoom with range of [0.94, 1.06], which means 94% zoom-in and 106% zoom-out, respectively. We generated 10,000 training images and 500 test images for each subset using data augmentation. The test images were augmented in order to have more images for testing the network model. Note that both the images and annotations were augmented in the same way.

We trained a state-of-the-art 2D-UNet network (Ronneberger et al. 2015) on the augmented training data using an ADAM optimizer with a learning rate of 0.0001. We used binary cross entropy as a loss function. The hyper parameter batch size was set to 2 and the step per epochs to 100. We trained the network for 50 epochs and assessed the quality of the models using the dice score. **Supplementary Figure 3a** compares dice score across all the subsets. We selected the model with the least percentage of split and dice score >80%. 20% of manual labeling was enough to predict the complete label fields for the forebrain, midbrain, retina, cerebellum and striatum periventricular, whereas telencephalon and neuropil required 25% and hindbrain 30%, respectively. Based on these results, the remaining 13 regions were then partially segmented by experts up to 20% and experiments were performed on these partially segmented regions to get the complete label fields.

### MECE and MISC masks

All of the zebrafish MECE masks and most of the MISC masks are available in the Z-brain atlas (https://fishexplorer.zib.de/zebrafish/lm/downloads/). The masks for serotonergic and oxytocin neurons were generated in the same way as the medaka MISC masks by using the Fiji plug-in, NeMO.

To compare the size of zebrafish and medaka brain region masks quantitatively, we visualized the relative volumetric data in a Manhattan plot. In order to organize the order of MECE masks, we sorted the brain regions from rostral to caudal direction, and assigned an individual number (region ID) to each brain MECE mask (total: 113).

For MISC masks, we assigned ID 000 - 099 to the MISC masks based on anatomy; ID 200 - 900 to MISC masks based on gene expression patterns. To have room for updating or adding new regions in the list, we assigned IDs for the gene expression masks starting with decimal numbers, for example 500.

### Brain registration

After the imaging step, all imaging volumes were registered to the reference brain using the Advanced Normalization Tools (ANTs) (Ahrens et al. 2012). We performed registration with rigid [-m GC, -c 300x300x300x0, 1e-8,10, -f 12x8x4x2, -s 4x3x2x1], affine [-m GC, -c 300x300x300x0, 1e-8,10, -f 12x8x4x2, -s 4x3x2x1] and SyN [-m CC, -c 300x300x300x0, 1e-8,10, -f 12x8x4x2x1, -s 4x3x2x1x0] with the parameters specified, respectively. The registrations were carried out on a high-performance cluster with 300 GB memory and 150 CPU cores. The tutorial with a detailed description of the parameters and step-by-step instructions for the registration with the Python script will be available on the webpage.

For the immunostained samples, we always stained with Hoechst dye, and used the Hoechst reference brain stacks for registration. To register HCR-stained samples with the reference brain, we had a few stacks of *snap25* and Hoechst-stained brains and made a *snap25* channel for the reference brain. HCR staining samples always had *snap25* channels to register other staining channels to the reference brain. After registration, we created the average stacks that were used to determine the region masks.

### cfos experiments and analysis

For the social stimuli assay, two fish larvae were placed at room temperature in a petri dish with light on. For the feeding assay, the fish larvae were placed in a dish and paramecia was added. For the MS222 assay, 60 ug/ml of MS222 (Sigma-Aldrich) were dissolved in filtered fish water. Samples were collected 40 minutes after treatments and immediately fixed with 4% PFA/PBS for overnight. For the vibration assay, we placed the dish with fish larvae on to the shaker (SCILOGEX, SCI-0180-S) at the speed of 60 rpm. Then HCR experiments were performed with *cfos* and *snap25* probes as described above. *Snap25* was used to normalize and quantify the *cfos* signal as it is broadly expressed in the nervous system.

To analyze the neural activity across samples in each treatment, we first created a snap25 channel for the reference brain. After registering the imaging stacks to the reference brain by aligning to this *snap25* channel, we calculated the median value of *cfos* and *snap25* staining across all MECE masks. The Manhattan plot shows the average values across the samples. All the values were normalized by the average values of the MECE masks. The average image stacks are generated by averaging each pixel across samples, then the maximum projected image is displayed. The lookup table was chosen so that the highest and lowest values matched the maximum and minimum of the lookup table. For the cfos analysis in the telencephalon, the same analysis was performed using the miscellaneous masks created for the telencephalon. Here, we normalized the cfos value of MISC region masks in the Manhattan plot to the value of the olfactory bulb in both species.

### Data access

The image data generated in this study for the medaka brain atlas will be submitted to Dryad. In addition, the image stacks of the reference brain and other gene channels are available in the FishExplorer (https://fishexplorer.zib.de/medaka/lm). The codes for analyzing brain region masks have been submitted to github. The tutorial with a detailed description of the parameters, step-by-step instructions for image registration and the Python script will be available on the webpage.

## Appendix medaka MECE regions

Appendix – medaka MECE regions
| ID | Name of MECE region |  | Methods | References | Notes |
| --- | --- | --- | --- | --- | --- |
| 1 | Whole brain |  |  |  |  |
| 2 | Eye |  | Hoechst, Ath5:GFP, HuC:DsRed |  | Different layers are labeled by three stainings. |
| 3 | Forebrain |  |  |  |  |
| 4 |  | Olfactory bulb | HuC:DsRed |  |  |
| 5 | Telencephalon |  |  |  |  |
| 6 |  | Pallium | HuC:DsRed, vglut:GFP, vglut (HCR) |  |  |
| 7 |  | Subpallium | Strong HuC:DsRed, gad1b (HCR) | (Okuyama et al. 2013) |  |
| 8 | Diencephalon |  |  |  |  |
| 9 |  | Pineal | Anti-5HT, Islet-1 (HCR) | (Lillesaar 2011) |  |
| 10 |  | Habenula | HuC:DsRed |  |  |
| 11 |  | Optic tract accessory, optic tectum | Ath5:GFP | (Robles, Laurell, and Baier 2014) | The boundary between this region and the stratum periventricular is not very clear. 5-HT positive signals around the anterior part of the diencephalon is used to set the boundaries of this region. |
| 12 |  | Arborization field 1 (AF1) | Ath5:GFP, Anti-Synapsin | (Robles, Laurell, and Baier 2014) |  |
| 13 |  | Arborization field 1 (AF2) | Ath5:GFP, Anti-Synapsin | (Robles, Laurell, and Baier 2014) |  |
| 14 |  | Arborization field 1 (AF3) | Ath5:GFP, Anti-Synapsin | (Robles, Laurell, and Baier 2014) |  |
| 15 |  | Arborization field 1 (AF4) | Ath5:GFP, Anti-Synapsin | (Robles, Laurell, and Baier 2014) |  |
| 16 |  | Arborization field 1 (AF5) | Ath5:GFP, Anti-Synapsin | (Robles, Laurell, and Baier 2014) |  |
| 17 |  | Arborization field 1 (AF6) | Ath5:GFP, Anti-Synapsin | (Robles, Laurell, and Baier 2014) |  |
| 18 |  | Arborization field 1 (AF7) | Ath5:GFP, Anti-Synapsin | (Robles, Laurell, and Baier 2014) | Outermost part of AF |
| 19 |  | Arborization field 1 (AF8) | Strong Anti-Synapsin | (Robles, Laurell, and Baier 2014) |  |
| 20 |  | Arborization field 1 (AF9) | Ath5:GFP, Anti-Synapsin | (Robles, Laurell, and Baier 2014) |  |
| 21 |  | Pretectum | Anti-5-HT, no HuC:DsRed, gad1b (HCR) cluster | (Antinucci, Folgueira, and Bianco 2019) (Lillesaar 2011) | Connect to AF7 in zebrafish, direct connect or next to AF |
| 22 |  | Thalamus | HuC:DsRed | (Mueller 2012) (Haehnel-Taguchi et al. 2018) |  |
| 23 |  | Dorsal thalamus | HuC positive cells in the rostral dorsal thalamus, non-HuC positive cells in the caudal dorsal thalamus. Gad1b negative | (Mueller 2012) |  |
| 24 |  | Ventral thalamus | HuC positive cells in the caudal ventral thalamus, Gad1b (HCR) positive | (Mueller 2012) |  |
| 25 |  | Posterior tuberculum | TH, 5-HT, HuC:DsRed | (Heap et al. 2018), (Schweitzer et al. 2012) (“Evolution of the Posterior Tuberculum and Preglomerular Nuclear Complex,” n.d.) (Lillesaar 2011) |  |
| 26 |  | Preglomerular complex | Presynapsin-positive signals, HuC:DsRed | (Bloch et al. 2020), (Wullimann and Umeasalugo 2020), (Kato, Yamada, and Yamamoto 2011) | The Preglomerular complex includes PGI, lateral preglomerular nucleus; PGM, medial preglomerular nucleus. |
| 27 |  | Preoptic area | TH, HuC:DsRed |  | Anterior to posterior tuberculum, posterior to subpallium |
| 28 |  | Rostral hypothalamus | HuC:DsRed, Anti-5-HT | (Lillesaar 2011) |  |
| 29 |  | Intermediate hypothalamus | HuC:DsRed, Hoechst, Anti-5 HT clusters in the middle | (Lillesaar 2011) | Ventral to the preglomerular complex. Area between the rostral and caudal hypothalamus. |
| 30 |  | Diffuse nucleus of the intermediate hypothalamus | Weak HuC:DsRed, Hoechst | (Heap et al. 2018) |  |
| 31 |  | Caudal hypothalamus | Ath5:GFP, Anti-5-HT | (Lillesaar 2011) |  |
| 32 |  | Pituitary | Anti-5HT |  |  |
| 33 | Midbrain |  |  |  |  |
| 34 |  | Torus Longitudinalis | Hoechst |  |  |
| 35 |  | Torus Semicircularis | Anti-synapsin, Anti-5-HT innervation, vglut(HCR) |  |  |
| 36 |  | Tectum |  |  |  |
| 37 |  | Tectum Neuropil | Projection from Ath5:GFP, no Hoechst signals |  |  |
| 38 |  | Tectm Stratum periventricular | Ath5:GFP, Hoechst |  |  |
| 39 |  | Tegmentum | Anti-TH (ventral area), Anti-ChAT cluster |  | Bridging between Torus Semicircular. Posterior to preglomerular complex |
| 40 |  | Tegmentum Oculomotor nucleus nlll | HuC:DsRed, vglut:GFP, islet-1(HCR) |  |  |
| 41 |  | Tegmentum NucMLF | Backfill | (Thiele, Donovan, and Baier 2014) |  |
| 42 | Hindbrain |  |  |  |  |
| 43 |  | Rhombomere 1 | Hoechst |  |  |
| 44 |  | Interpeduncular nucleus | HuC:DsRed |  | Rhombomere 1 |
| 45 |  | Raphe superior | Anti-5-HT | (Lillesaar 2011) | Rhombomere 1, 2 |
| 46 |  | Oculomotor nucleus | vglut:GFP, islet-1 (HCR) |  | Rhombomere 1, cranial nerve |
| 47 |  | R1_RoM1 | Backfill |  |  |
| 48 |  | R123_RoL | Backfill |  |  |
| 49 |  | Locus Coeruleus | Anti-TH |  | Rhombomere 1,2<br>Noradrenergic neurons labeled by Anti-TH |
| 50 |  | Cerebellum | HuC:DsRed |  |  |
| 51 |  | Cerebellum Cerebellar Corpus |  |  |  |
| 52 |  | Cerebellum Valvula Cerebelli | Anti-5HT innervation | (Lillesaar 2011) |  |
| 53 |  | Cerebellum Medial caudal lobe | HuC:DsRed, Hoechst |  |  |
| 54 |  | Cerebellum Granular Eminence | HuC:DsRed |  |  |
| 55 |  | Rhombomere 2 | Hoechst |  |  |
| 56 |  | Anterior_Cluster_of_nV_Trigenimal_motor_neruons | Islet-1 (HCR) | (Higashijima, Hotta, and Okamoto 2000) |  |
| 57 |  | Raphe superior | Anti-5-HT | (Lillesaar 2011) | Rhombomere 1, 2 |
| 58 |  | R123_RoL | Backfill |  |  |
| 59 |  | R2_RoM2 | Backfill |  |  |
| 60 |  | Locus Coeruleus | Anti-TH |  | Rhombomere 1,2<br>Noradrenergic neurons labeled by<br>Anti-TH |
| 61 |  | Rhombomere 3 | Hoechst |  |  |
| 62 |  | Posterior_Cluster_of_nV_Trigeminal_motor_neurons | Islet-1 (HCR) | (Higashijima, Hotta, and Okamoto 2000) |  |
| 63 |  | R3_RoM3 | Backfill |  |  |
| 64 |  | R123_RoL | Backfill |  |  |
| 65 |  | RoV3 |  |  | RoV3 doesn't exist in medaka |
| 66 |  | Raphe inferior | Anti-5-HT |  | Rhombomere 3,4,5,6,7 |
| 67 |  | Rhombomere 4 | Hoechst |  | Mauthner cell locates in<br>Rhombomere 4 |
| 68 |  | R4_Mauthner |  |  |  |
| 69 |  | Mauthner_cell_Axon_Cap | Anti-synapsin | (Bierman, Zottoli, and Hale 2009) |  |
| 70 |  | Raphe inferior | Anti-5-HT |  | Rhombomere 3,4,5,6,7 |
| 71 |  | R4_MiV1 | Backfill |  |  |
| 72 |  | R4_MiM1 | Backfill |  |  |
| 73 |  | MiR1 |  |  | Medaka doesn't have MiR1. |
| 74 |  | MiR2 |  |  | Medaka doesn't have MiR2. |
| 75 |  | R3456_Vestibular | Backfill |  |  |
| 76 |  | Rhombomere 5 | Hoechst |  |  |
| 77 |  | R3456_Vestibular | Backfill |  |  |
| 78 |  | R5_MiD2 | Backfill |  |  |
| 79 |  | R5_MiV2 | Backfill |  |  |
| 80 |  | R5_MiR2 | Backfill |  | Zebrafish doesn't have MiR2 |
| 81 |  | Tangential_Vestibular_nucleus |  |  |  |
| 82 |  | VII_Facial_Motor_and_octavolateralis_efferent_neuron | Islet-1 (HCR) | (Higashijima, Hotta, and Okamoto 2000) |  |
| 83 |  | Rhombomere 6 | Hoechst |  |  |
| 84 |  | R3456_Vestibular | Backfill |  |  |
| 85 |  | R6_MiD3 | Backfill |  |  |
| 86 |  | R6_MiT | Backfill |  |  |
| 87 |  | Raphe inferior | Anti-5-HT |  | Rhombomere 3,4,5,6,7 |
| 88 |  | Tangential_Vestibular_nucleus | Backfill |  |  |
| 89 |  | VII_Facial_Motor_and_octavolateralis_efferent_neuron | Islet-1 (HCR) | (Higashijima, Hotta, and Okamoto 2000) |  |
| 90 |  | Rhombomere 7 | Hoechst |  |  |
| 91 |  | R7_CaD | Backfill |  |  |
| 92 |  | R7_CaV | Backfill |  |  |
| 93 |  | Raphe inferior | Anti-5-HT |  | Rhombomere 3,4,5,6,7 |
| 94 |  | VII_Facial_Motor_and_octavolateralis_efferent_neuron | Islet-1 (HCR) | (Higashijima, Hotta, and Okamoto 2000) |  |
| 95 |  | Caudal Hindbrain |  |  |  |
| 96 |  | Caudal Hindbrain<br>Area postrema | Anti-TH |  |  |
| 97 |  | Caudal Hindbrain<br>Lateral reticular nucleus | Anti-TH |  |  |
| 98 |  | Caudal Hindbrain<br>Inferior Olive | HuC:DsRed | (Felix et al., n.d.) |  |
| 99 |  | Caudal Hindbrain<br>X-vagus motor neuron cluster | Anti-TH, islet-1 (HCR) |  | Cranial nerve (X) |
| 100 | Spinal cord |  | HuC:DsRed, Anti-5 HT cluster, Anti-TH, HCR-ChAT |  | vglut:GFP positive signals in muscles |
| 101 | Ganglia |  |  |  |  |
| 102 |  | Olfactory epithelium | vglut:GFP, Hoechst, Ath5:GFP |  |  |
| 103 |  | Facial sensory ganglion | HuC:DsRed |  |  |
| 104 |  | Facial Glossopharyngeal ganglion (gIX) | Hoechst, islet-1(HCR) | (Cox et al. 2011) | Connected to Cranial nerve (IX) |
| 105 |  | Statoacoustic ganglion (VIII) | Tg (vglut:GFP) |  | The sensory ganglion of the ear. Connected to Cranial nerve (VIII) |
| 106 |  | Trigeminal ganglion | Ath5:GFP, Hoechst |  | Connected to Cranial nerve (V) |
| 107 |  | Anterior lateral line ganglion | Ath5:GFP, Hoechst |  |  |
| 108 |  | *Posterior lateral line ganglion* | Hoechst |  |  |
| 109 |  | Vagus ganglion (gX) | Hoechst, islet-1(HCR) | (Higashijima, Hotta, and Okamoto 2000) | Connected to Cranial nerve (X) |
| 110 |  | Lateral_line_Neuromast_D1 | FM1-34, anti-TH | (Haehnel-Taguchi et al. 2018; Manuel et al. 2021) |  |
| 111 |  | Lateral_line_Neuromast_D2 | FM1-34, anti-TH |  |  |
| 112 |  | Lateral_line_Neuromast_N | FM1-34, anti-TH |  |  |
| 113 |  | Lateral_line_Neuromast_O1 | FM1-34, anti-TH |  |  |
| 114 |  | Lateral_line_Neuromast_OC1 | FM1-34, anti-TH |  |  |
| 115 |  | Lateral_line_Neuromast_SO1 | FM1-34, anti-TH |  |  |
| 116 |  | *Lateral_line_Neuromast_SO2* | FM1-34, anti-TH |  |  |
| 117 |  | Lateral_line_Neuromast_SO3 | FM1-34, anti-TH |  |  |

## Appendix medaka MISC regions

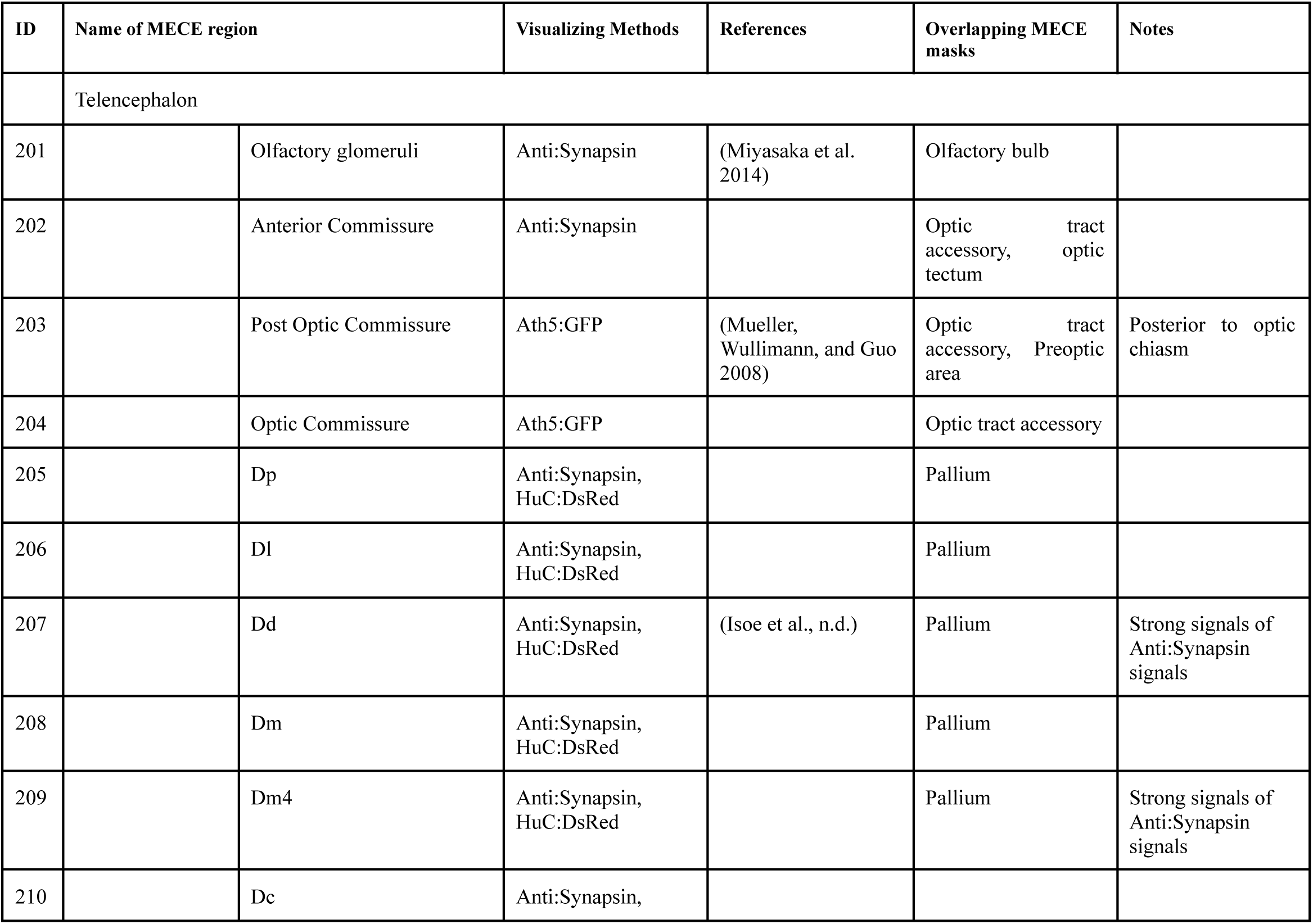
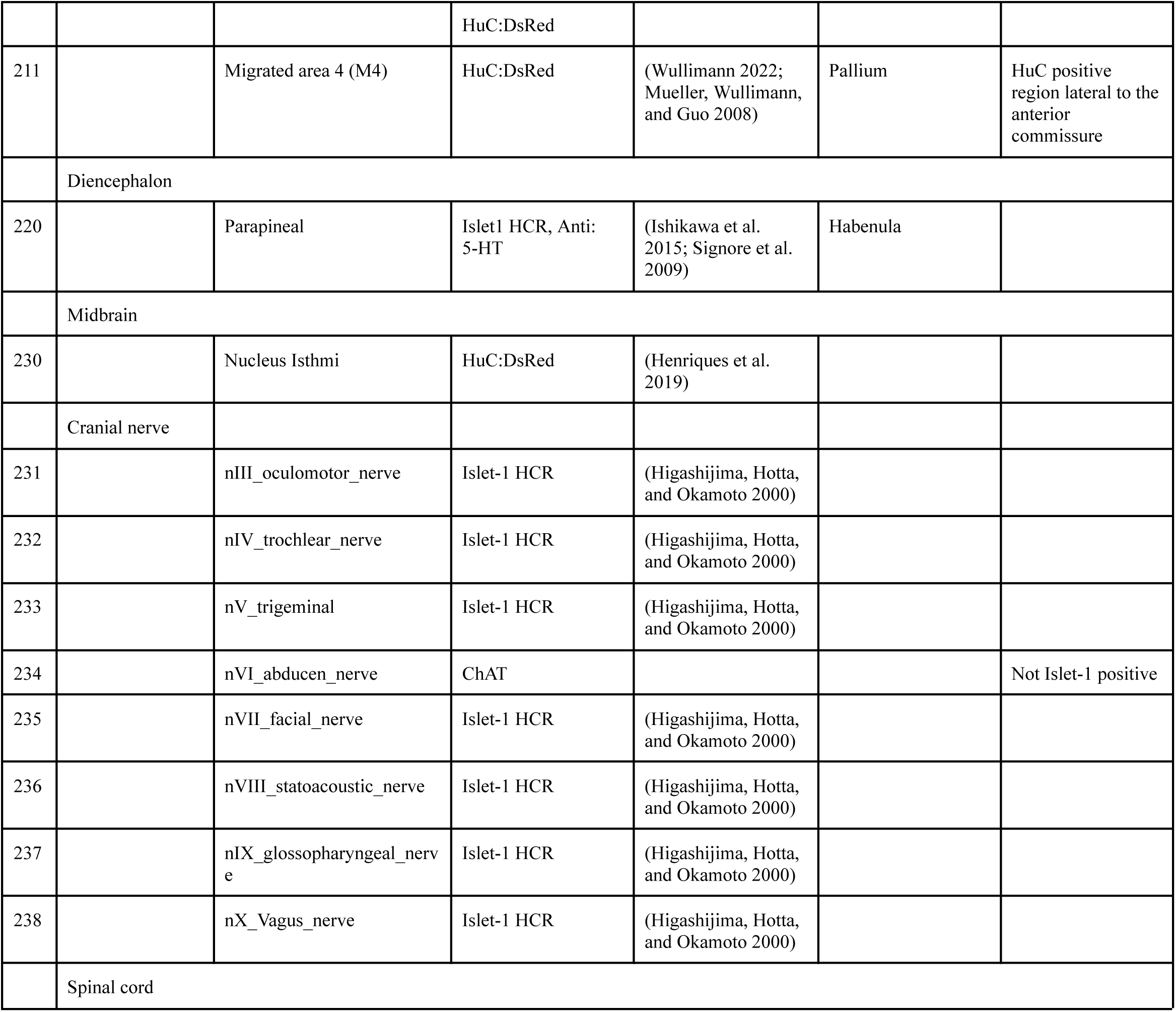

## Appendix medaka MISC region-gene clusters (neurotransmitters, neuropeptides)

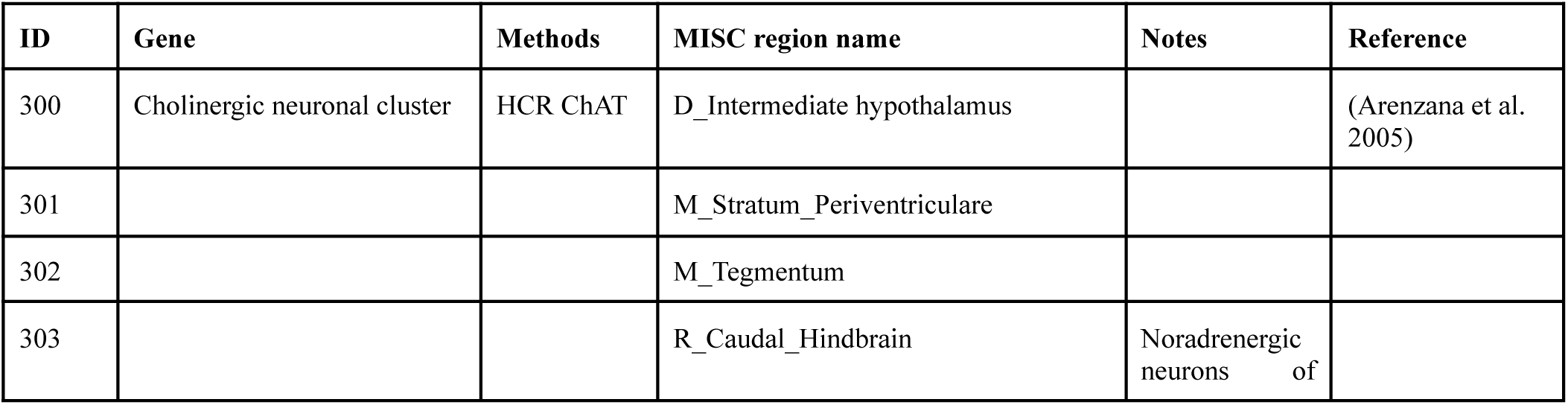
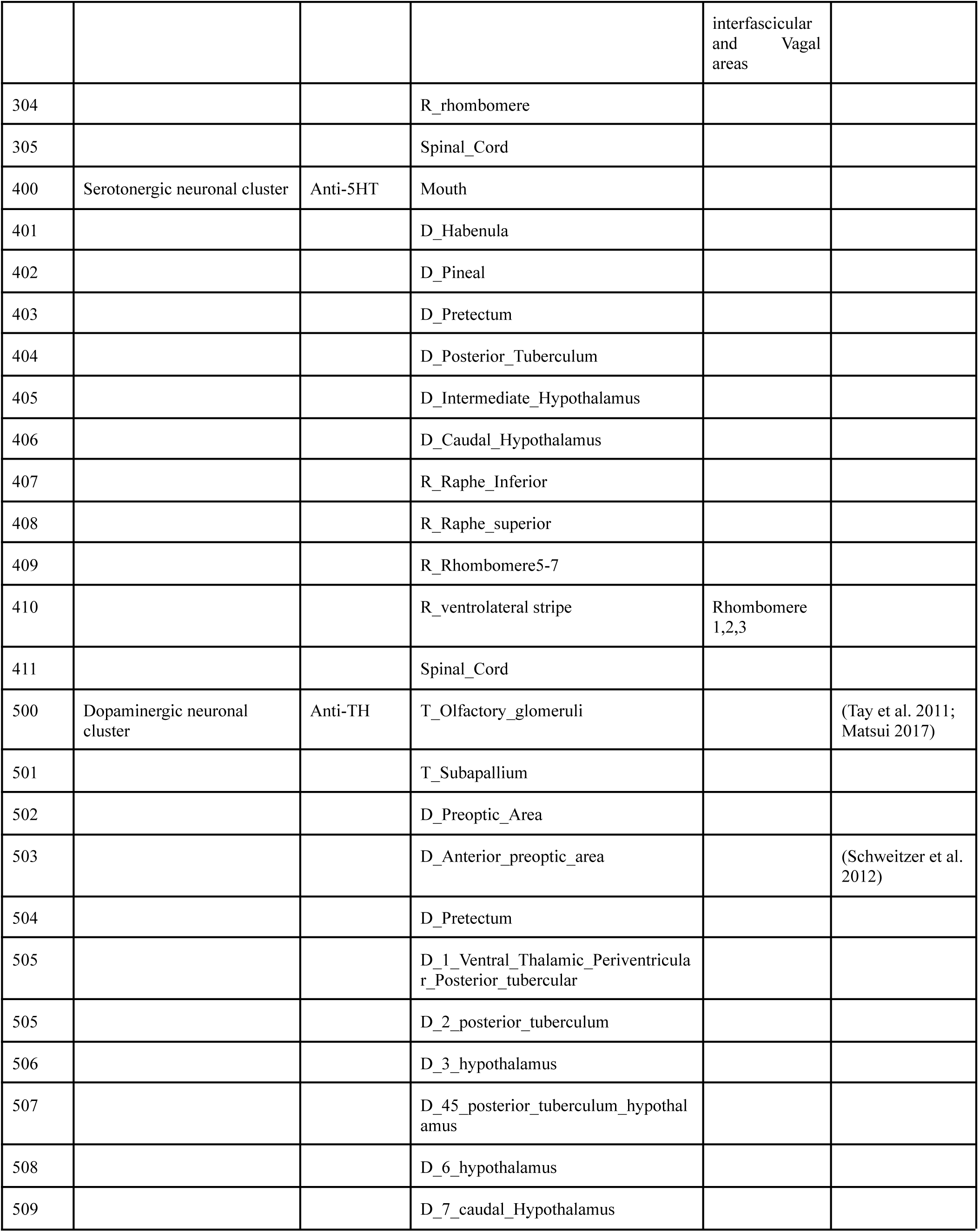
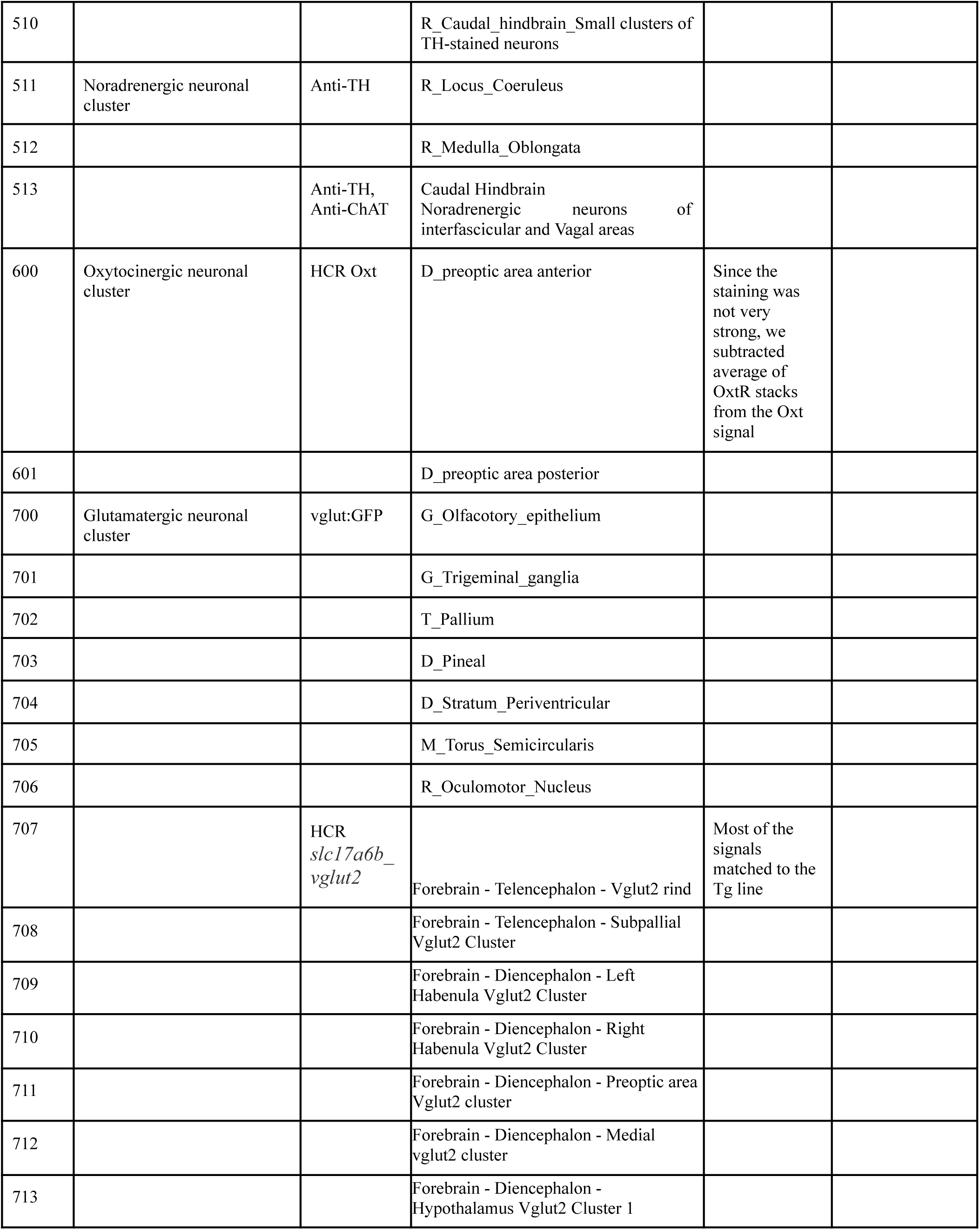
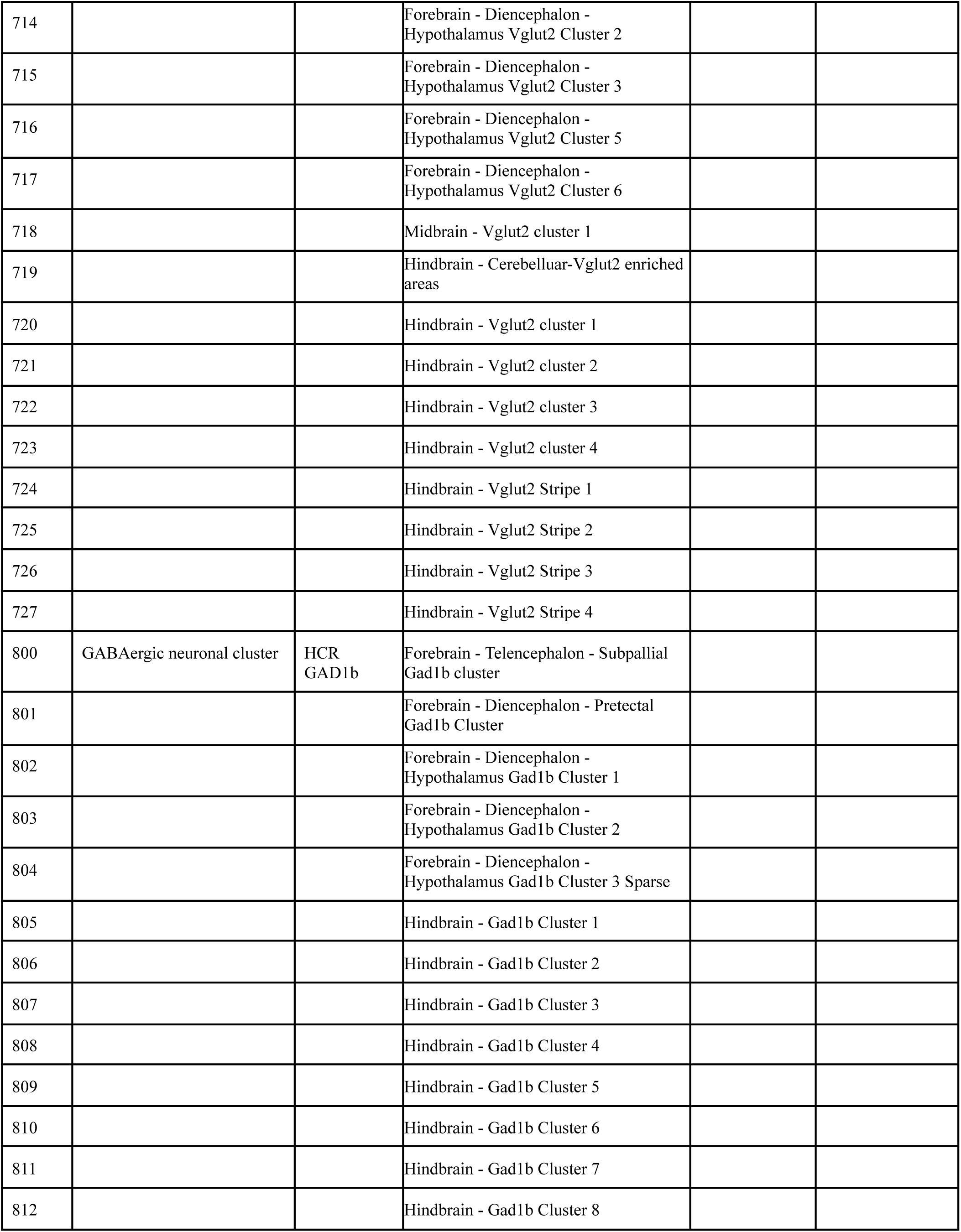
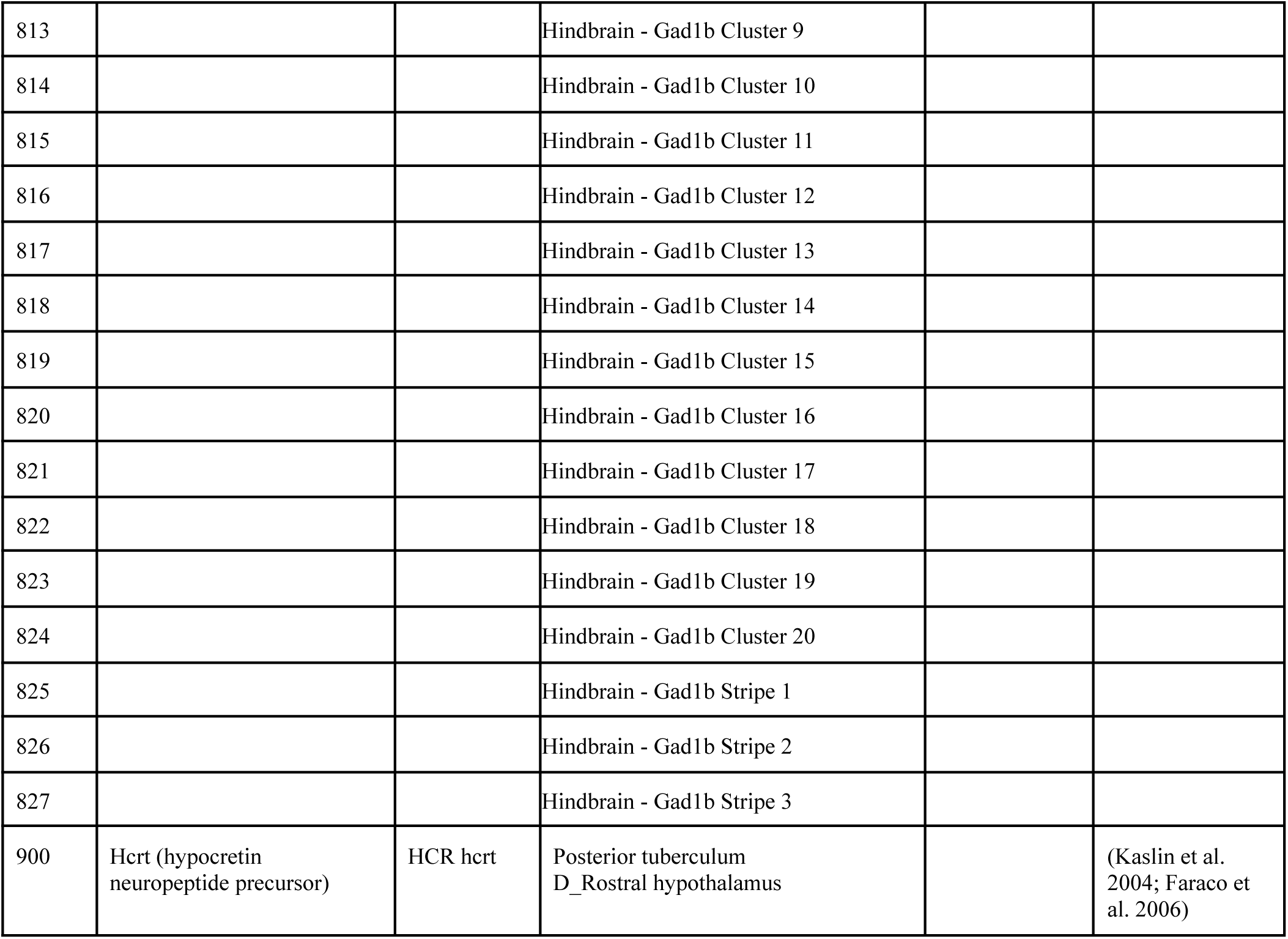

Appendix - medaka MISC region- gene clusters (neurotransmitters, neuropeptides)
| ID | Gene | Methods | MISC region name | Notes | Reference |
| --- | --- | --- | --- | --- | --- |
| 300 | Cholinergic neuronal cluster | HCR ChAT | D_Intermediate hypothalamus |  | (Arenzana et al. 2005) |
| 301 |  |  | M_Stratum_Periventriculare |  |  |
| 302 |  |  | M_Tegmentum |  |  |
| 303 |  |  | R_Caudal_Hindbrain | Noradrenergic neurons of interfascicular<br>and Vagal<br>areas |  |
| 304 |  |  | R_rhombomere |  |  |
| 305 |  |  | Spinal_Cord |  |  |
| 400 | Serotonergic neuronal cluster | Anti-5HT | Mouth |  |  |
| 401 |  |  | D_Habenula |  |  |
| 402 |  |  | D_Pineal |  |  |
| 403 |  |  | D_Pretectum |  |  |
| 404 |  |  | D_Posterior_Tuberculum |  |  |
| 405 |  |  | D_Intermediate_Hypothalamus |  |  |
| 406 |  |  | D_Caudal_Hypothalamus |  |  |
| 407 |  |  | R_Raphe_Inferior |  |  |
| 408 |  |  | R_Raphe_superior |  |  |
| 409 |  |  | R_Rhombomere5-7 |  |  |
| 410 |  |  | R_ventrolateral stripe | Rhombomere<br>1,2,3 |  |
| 411 |  |  | Spinal_Cord |  |  |
| 500 | Dopaminergic neuronal cluster | Anti-TH | T_Olfactory_glomeruli |  | (Tay et al. 2011;<br>Matsui 2017) |
| 501 |  |  | T_Subapallium |  |  |
| 502 |  |  | D_Preoptic_Area |  |  |
| 503 |  |  | D_Anterior_preoptic_area |  | (Schweitzer et al.<br>2012) |
| 504 |  |  | D_Pretectum |  |  |
| 505 |  |  | D_1_Ventral_Thalamic_Periventricular<br>r_Posterior_tubercular |  |  |
| 505 |  |  | D_2_posterior_tuberculum |  |  |
| 506 |  |  | D_3_hypothalamus |  |  |
| 507 |  |  | D_45_posterior_tuberculum_hypothalamus |  |  |
| 508 |  |  | D_6_hypothalamus |  |  |
| 509 |  |  | D_7_caudal_Hypothalamus |  |  |
| 510 |  |  | R_Caudal_hindbrain_Small clusters of TH-stained neurons |  |  |
| 511 | Noradrenergic neuronal cluster | Anti-TH | R_Locus_Coeruleus |  |  |
| 512 |  |  | R_Medulla_Oblongata |  |  |
| 513 |  | Anti-TH, Anti-ChAT | Caudal Hindbrain Noradrenergic neurons of interfascicular and Vagal areas |  |  |
| 600 | Oxytocinergic neuronal cluster | HCR Oxt | D_preoptic area anterior | Since the staining was not very strong, we subtracted average of OxtR stacks from the Oxt signal |  |
| 601 |  |  | D_preoptic area posterior |  |  |
| 700 | Glutamatergic neuronal cluster | vglut:GFP | G_Olfactory_epithelium |  |  |
| 701 |  |  | G_Trigeminal_ganglia |  |  |
| 702 |  |  | T_Pallium |  |  |
| 703 |  |  | D_Pineal |  |  |
| 704 |  |  | D_Stratum_Periventricular |  |  |
| 705 |  |  | M_Torus_Semicircularis |  |  |
| 706 |  |  | R_Oculomotor_Nucleus |  |  |
| 707 |  | HCR *slc17a6b_vglut2* | Forebrain - Telencephalon - Vglut2 rind | Most of the signals matched to the Tg line |  |
| 708 |  |  | Forebrain - Telencephalon - Subpallial Vglut2 Cluster |  |  |
| 709 |  |  | Forebrain - Diencephalon - Left Habenula Vglut2 Cluster |  |  |
| 710 |  |  | Forebrain - Diencephalon - Right Habenula Vglut2 Cluster |  |  |
| 711 |  |  | Forebrain - Diencephalon - Preoptic area Vglut2 cluster |  |  |
| 712 |  |  | Forebrain - Diencephalon - Medial vglut2 cluster |  |  |
| 713 |  |  | Forebrain - Diencephalon - Hypothalamus Vglut2 Cluster 1 |  |  |
| 714 |  |  | Forebrain - Diencephalon - Hypothalamus Vglut2 Cluster 2 |  |  |
| 715 |  |  | Forebrain - Diencephalon - Hypothalamus Vglut2 Cluster 3 |  |  |
| 716 |  |  | Forebrain - Diencephalon - Hypothalamus Vglut2 Cluster 5 |  |  |
| 717 |  |  | Forebrain - Diencephalon - Hypothalamus Vglut2 Cluster 6 |  |  |
| 718 |  |  | Midbrain - Vglut2 cluster 1 |  |  |
| 719 |  |  | Hindbrain - Cerebellar-Vglut2 enriched areas |  |  |
| 720 |  |  | Hindbrain - Vglut2 cluster 1 |  |  |
| 721 |  |  | Hindbrain - Vglut2 cluster 2 |  |  |
| 722 |  |  | Hindbrain - Vglut2 cluster 3 |  |  |
| 723 |  |  | Hindbrain - Vglut2 cluster 4 |  |  |
| 724 |  |  | Hindbrain - Vglut2 Stripe 1 |  |  |
| 725 |  |  | Hindbrain - Vglut2 Stripe 2 |  |  |
| 726 |  |  | Hindbrain - Vglut2 Stripe 3 |  |  |
| 727 |  |  | Hindbrain - Vglut2 Stripe 4 |  |  |
| 800 | GABAergic neuronal cluster | HCR GAD1b | Forebrain - Telencephalon - Subpallial Gad1b cluster |  |  |
| 801 |  |  | Forebrain - Diencephalon - Pretectal Gad1b Cluster |  |  |
| 802 |  |  | Forebrain - Diencephalon - Hypothalamus Gad1b Cluster 1 |  |  |
| 803 |  |  | Forebrain - Diencephalon - Hypothalamus Gad1b Cluster 2 |  |  |
| 804 |  |  | Forebrain - Diencephalon - Hypothalamus Gad1b Cluster 3 Sparse |  |  |
| 805 |  |  | Hindbrain - Gad1b Cluster 1 |  |  |
| 806 |  |  | Hindbrain - Gad1b Cluster 2 |  |  |
| 807 |  |  | Hindbrain - Gad1b Cluster 3 |  |  |
| 808 |  |  | Hindbrain - Gad1b Cluster 4 |  |  |
| 809 |  |  | Hindbrain - Gad1b Cluster 5 |  |  |
| 810 |  |  | Hindbrain - Gad1b Cluster 6 |  |  |
| 811 |  |  | Hindbrain - Gad1b Cluster 7 |  |  |
| 812 |  |  | Hindbrain - Gad1b Cluster 8 |  |  |
| 813 |  |  | Hindbrain - Gad1b Cluster 9 |  |  |
| 814 |  |  | Hindbrain - Gad1b Cluster 10 |  |  |
| 815 |  |  | Hindbrain - Gad1b Cluster 11 |  |  |
| 816 |  |  | Hindbrain - Gad1b Cluster 12 |  |  |
| 817 |  |  | Hindbrain - Gad1b Cluster 13 |  |  |
| 818 |  |  | Hindbrain - Gad1b Cluster 14 |  |  |
| 819 |  |  | Hindbrain - Gad1b Cluster 15 |  |  |
| 820 |  |  | Hindbrain - Gad1b Cluster 16 |  |  |
| 821 |  |  | Hindbrain - Gad1b Cluster 17 |  |  |
| 822 |  |  | Hindbrain - Gad1b Cluster 18 |  |  |
| 823 |  |  | Hindbrain - Gad1b Cluster 19 |  |  |
| 824 |  |  | Hindbrain - Gad1b Cluster 20 |  |  |
| 825 |  |  | Hindbrain - Gad1b Stripe 1 |  |  |
| 826 |  |  | Hindbrain - Gad1b Stripe 2 |  |  |
| 827 |  |  | Hindbrain - Gad1b Stripe 3 |  |  |
| 900 | Hcrt (hypocretin<br>neuropeptide precursor) | HCR hcrt | Posterior tuberculum<br>D_Rostral hypothalamus |  | (Kaslin et al.<br>2004; Faraco et<br>al. 2006) |

## Authors contributions

Conceptualization: SV, FE, YI, Analysis: SV, YI, Software: SV, Visualization: SV, TTS, YI, Writing original draft: SV, YI, Writing review & editing: SV, TTS, HCH, DB, FE, YI, Methodology: SV, KH, TTS, JBW, SK, AS, SC, CS,YI, Resources: JW, AA, MF, FE, Experimental support: MF, FE, Funding acquisition: MF, FE, Supervision: MF, HCH, DB, FE

## Acknowledgements

We are grateful to the Engert lab and the Fishman lab in Harvard University for experimental and thoughtful support. We appreciate the Bellono lab in Harvard University for experimental support in making transgenic lines. We acknowledge Epoch for experimental support in making transgenic lines. We thank the Armin Bahl lab in Konstanz Institute for experimental support and kind suggestions in brain registration. We appreciate the Harvard Center for Biological Imaging (HCBI) for microscopic imaging. We acknowledge Iris Odstrcil for the help in backfill experiments. We appreciate Erin Yue Song for HCR experiments. We thank the Shinichi Higashijima lab in NIBB for sharing the Tg (Glut:GFP) line.

---

**Supplementary Figure 1:**
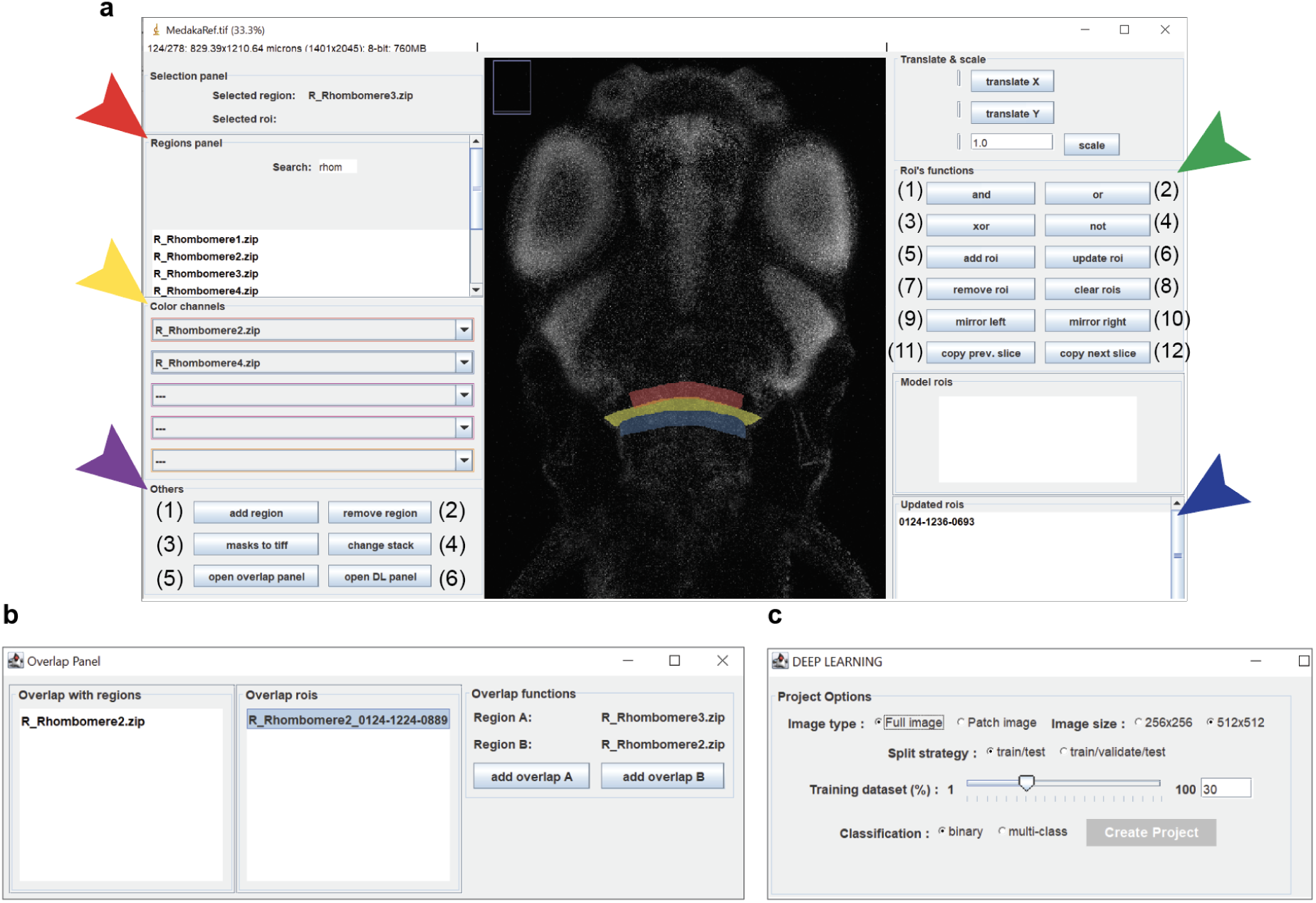
Medaka reference brain. Reference brain stacks. We have three channels in the medaka reference brain: (1) tERK channel: total extracellular signal-regulated kinase (tERK) is stained in the whole cell bodies, which is used for Z-brain reference brain and used to register neural activity with phospho-ERK (pERK)/tERK staining (Randlett et al. 2015); (2) Hoechst channel: All nuclei are labeled by Hoechst staining (Latt et al. 1975); (3) HuC promoter channel: The HuC (*elav*) promoter is driven pan-neuronally and is commonly used for GCaMP-based calcium imaging for analyzing *in vivo* neural activity (Ahrens et al. 2012). The larvae of Tg (HuC:DsRed, red) line were stained with anti-tERK immunohistochemistry (green) and Hoechst dye (blue).

**Supplementary Figure 2:**
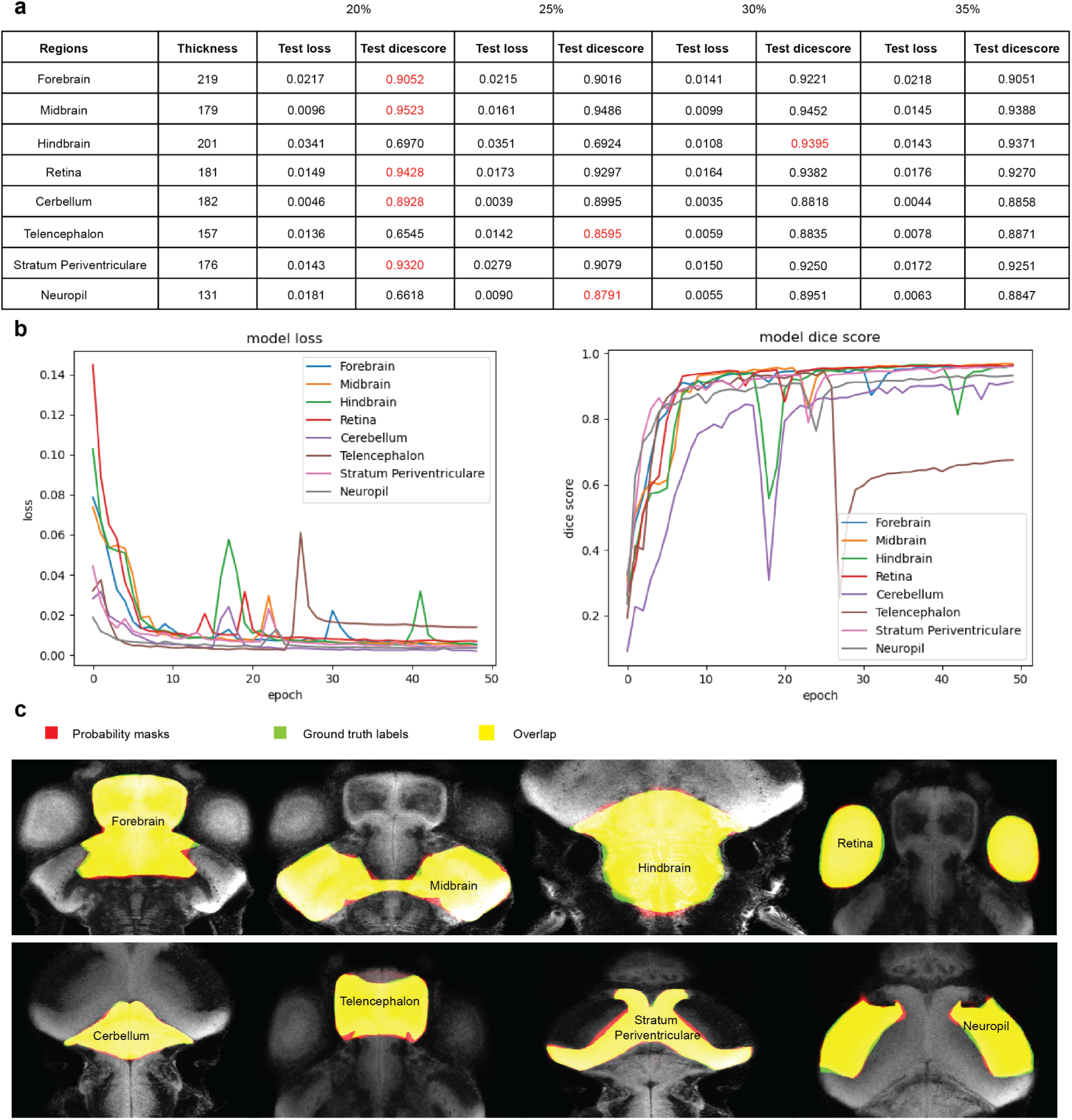
A new interactive Fiji tool - Neuro Mask Organizer (NeMO) - for editing masks. (a) The Fiji plug-in has various utility functions, including the comparison of masks with registered regions and fixing overlaps between regions. By visualizing many adjacent region masks, new masks can be drawn and added. The red arrowhead shows the list of the masks to be modified, the yellow arrowhead shows the list of masks in different colors. The purple arrowhead shows the function buttons: (1) “add region”, (2) “remove region”, (3) “masks to tiff”, (4) “change stack”, (5) “open overlap panel”, and (6) “open DL panel”. The green arrowhead indicates the functions: (1) “and”, (2) “or”, (3) “xor”, (4) “not”, (5) “add roi”, (6) “update roi”, (7) “remove roi”, (8) “clear rois”, (9) “mirror left”, (10) “mirror right”, (11) “copy prev. slice”, (12) “copy next slice”. And the blue arrowhead indicates regions of interest (ROIs) that are displayed in yellow. (b) The overlap panel provides utility functions to fix overlaps between regions. (c) The DL panel allows users to create a training dataset based on different parameters, including classification strategy (binary/multiclass), split strategy (train/test vs. train/validate/test), and training on full images or patches. The parameters chosen and the rationale behind are described in the Methods section.

**Supplementary Figure 3:**
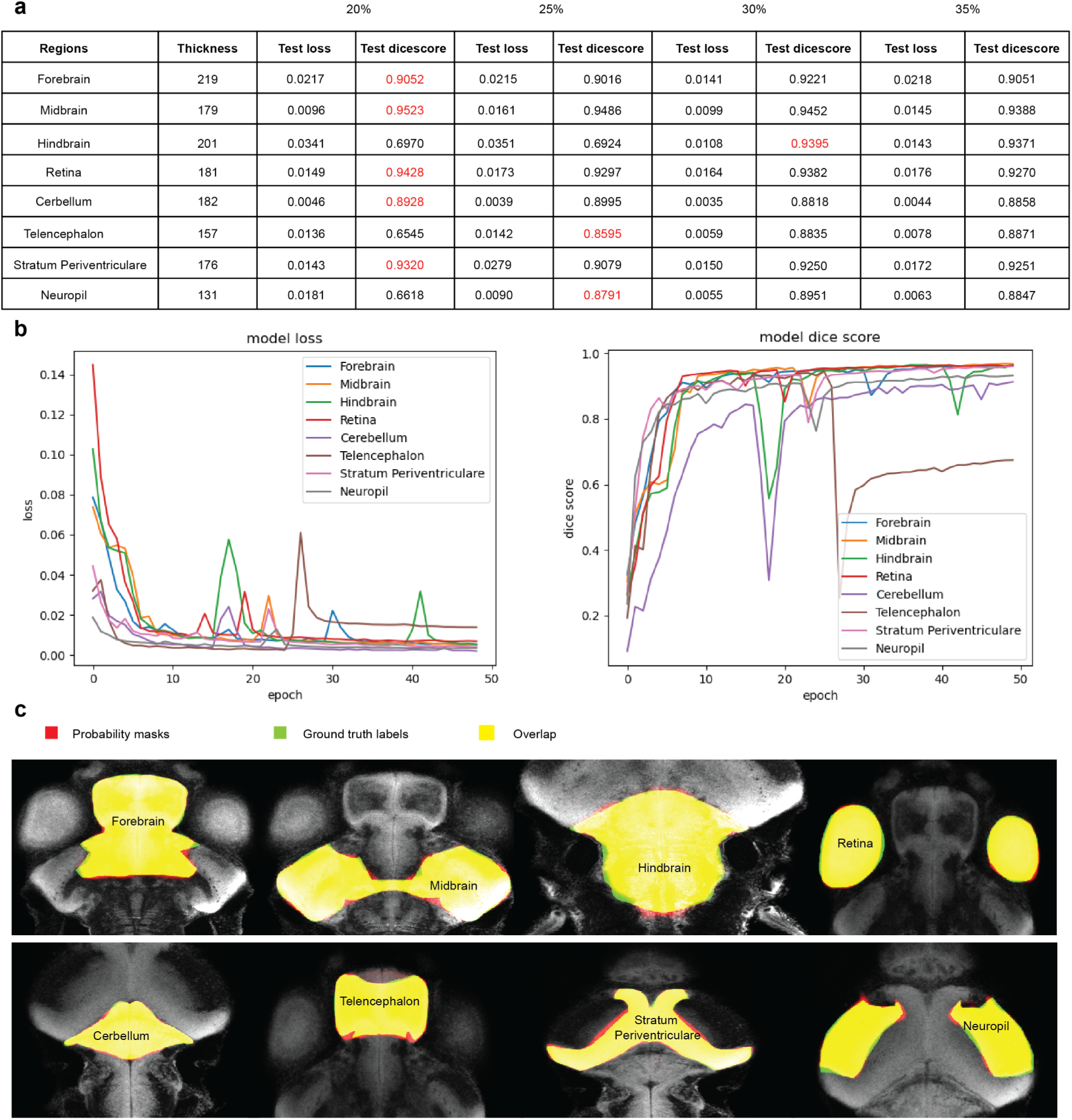
Segmentation of brain regions using deep learning. (a) Comparison of segmentation results for binary classification of 8 regions with 20%, 25%, 30%, and 35% of images for training, respectively. For each region, a model with the least training data and a dice coefficient above 80% is selected (marked as red). (b) Training curves of binary cross entropy losses and dice scores for the selected models. (c) The slice views show the classifier prediction (red), ground truth (green) and overlap (yellow) for each region using the selected model.

**Supplementary Figure 4:**
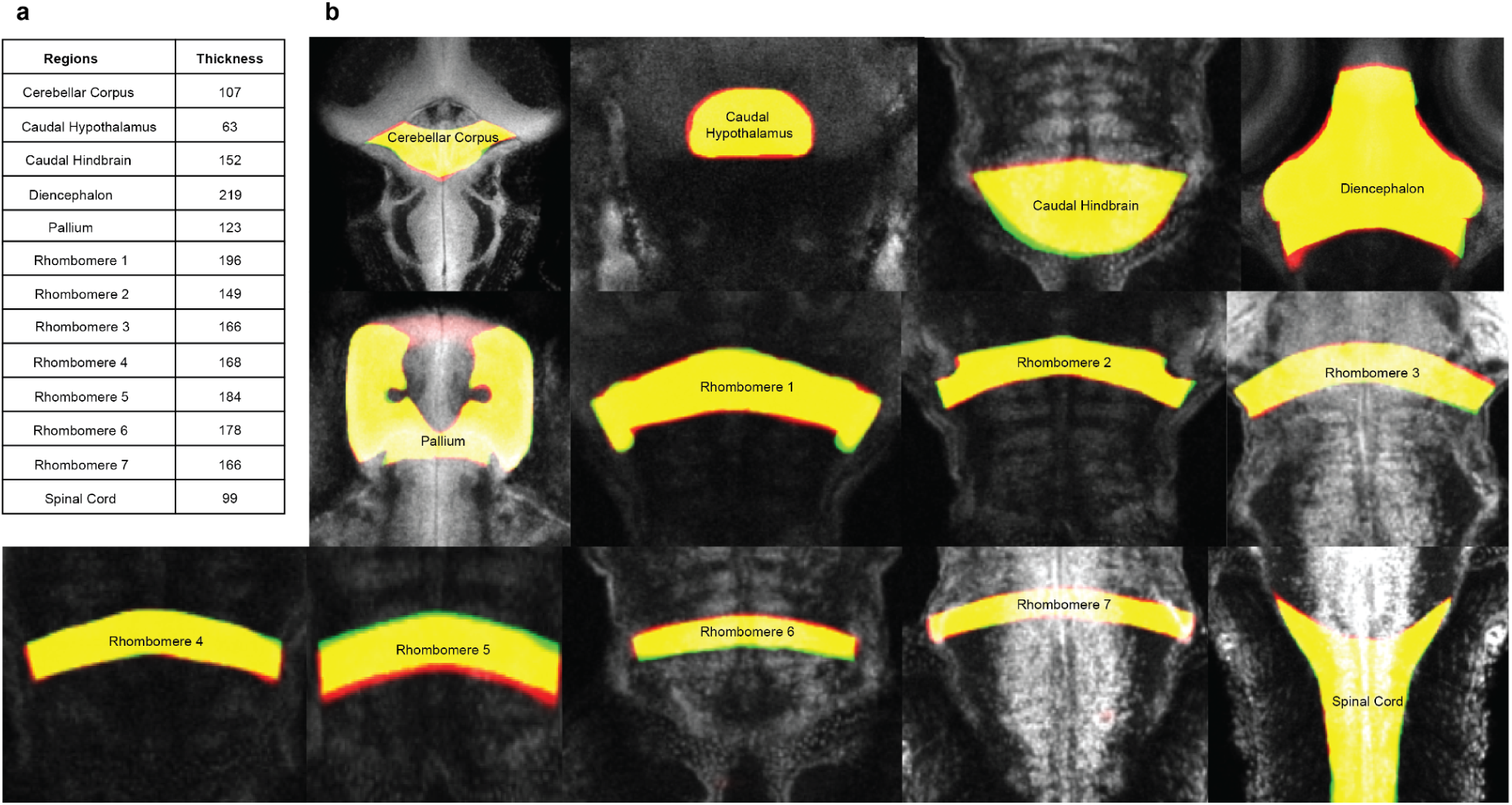
Segmentation of brain regions using deep learning. (a) List of 13 regions and their respective thickness; (b) The slice views show the classifier prediction (red), ground truth (green) and overlap (yellow) for each region.

**Supplementary Figure 5:**
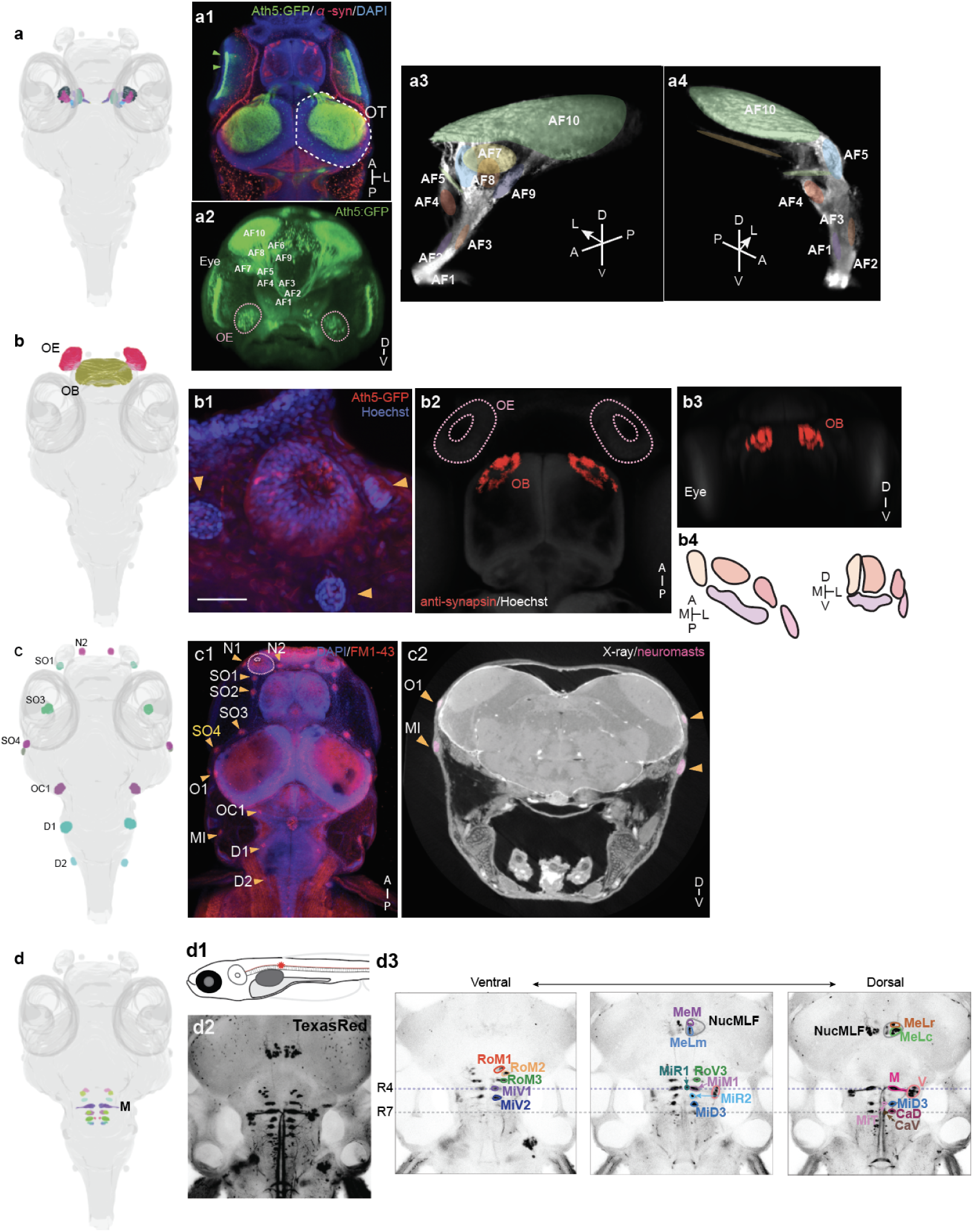
MISC masks of sensorimotor pathways. (a) <u>Retinotectal projection.</u> (a1) The Tg (Ath5:GFP) line marks the axonal projection and arborization fields (AFs) of the retinal ganglion cells. The anti-synapsin immunostaining onto the Tg line allowed us to categorize the arborization field (left). OT: optic tectum. Green triangles indicate the retinal ganglion cells labeled in the Tg (Ath5:GFP) line. A: anterior axis, P: posterior axis, and L: lateral direction. (a2) A view from the head of the 3D reconstruction of the confocal imaging of larvae of the Tg (Ath5:GFP) line. Based on the location and axonal projections, we identified ten arborization fields (AF). In the Tg line, not only retinal ganglion projections but also cells in the olfactory epithelium (OE) are visualized (pink dotted line circles). D-V indicates the dorso-ventral axis. (a3, 4) Arborization fields (AF1-9 and Neuropil (AF10)). Lateral view from outside (a3) and from inside (a4). (b) <u>Olfaction.</u> The olfactory epithelium (OE) (b1) and the olfactory bulb (OB) (b2 - 4). (b1) OE is visualized by Tg (Ath5-GFP) and Hoechst staining (blue: Hoechst, red: Ath5-GFP). OE is surrounded by neuromasts (orange triangles). (b2) Dorsal (b2) and front view (b3) of OB, visualized by anti-synapsin immunostaining (red). Neurons in OE project into five olfactory glomeruli in each hemisphere of OB (b4). (c) <u>Visualization of neuromasts.</u> (c1) Neuromasts are visualized by staining with FM1-43 membrane probes. The dorsal view is shown. Orange triangles indicate the position of neuromasts around the head of the medaka larva. Medaka-specific neuromasts SO4 are highlighted in yellow. N: Nasal neuromasts, SO: supraorbital neuromasts, O: optic neuromasts, OC: occipital neuromasts, MI: middle neuromasts, D: dorsal neuromasts. (c2) X-ray imaging shows the protruding structure of neuromasts (pink). (d) <u>Reticulospinal projection.</u> Backfill injection was done in the floor plate of the spinal cord (d1) to visualize the reticulospinal projection in the midbrain and hindbrain (dorsal view, d2). We identified cellular clusters in the reticulospinal cord (d3). Backfill-visualized neurons in medaka larvae are shown. The visualized cell clusters were identified by their relative position to the Mauthner cells (M), which are identified by their unique structure. The top dotted line indicates the same position to the Mauthner cells in the AP axis (in the rhombomere 4, R4), and the bottom dotted line indicates the same position to the most posterior cell in the middle area (in the rhombomere 7, R7). M (Mauthner cell), RoM (rostral of Mauthner cells) clusters, MiM (middle cells in middle area), MiV (ventral cells in middle area), MiD (dorsal cells in middle area), MiT (T-shaped axon in middle area), MiR (rostral cells in middle area), MeM (projection in mesencephalon, medial), MeLm (projection in mesencephalon, lateral medial), MeLr (projection in mesencephalon, rostral), MeLc (projection in mesencephalon, caudal), RoV3 (projection in rhombencephalon, ventral), CaD (dorsal cells in caudal area), CaV (ventral cells in caudal area), nucMLF (nucleus of medial longitudinal fasciculus), V (vestibular clusters) and its axonal projection was visualized.

**Supplementary Figure 6:**
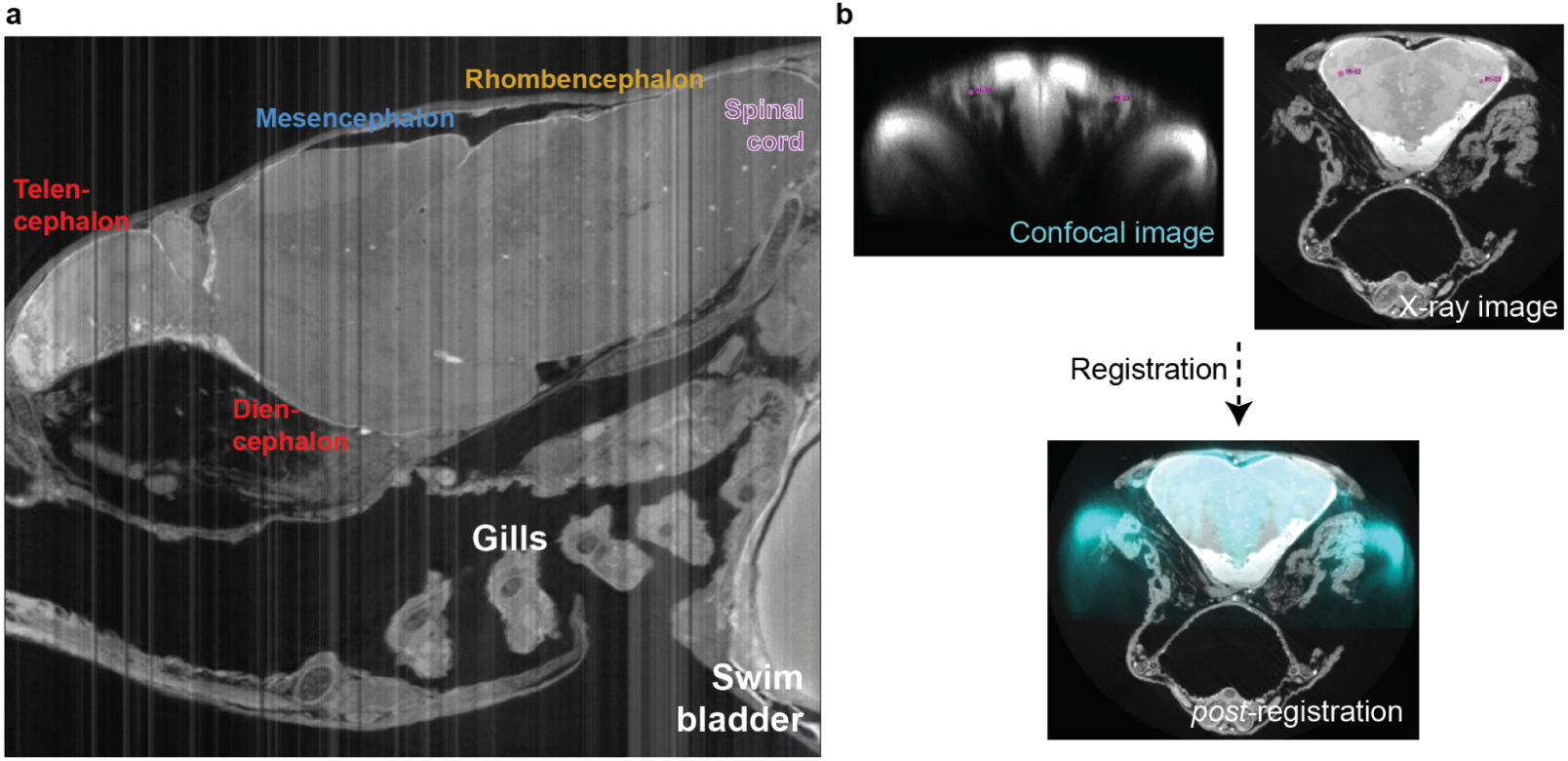
X-ray imaging of medaka larvae. (a) Lateral view of a medaka larva by X-ray. The telencephalon, diencephalon, mesencephalon, rhombencephalon and spinal cord are labeled. With X-ray imaging we also observed the gills and swim bladder. (b) The confocal imaging stack was registered onto the X-ray image stack to align two imaging stacks from both modalities.

